# Global RNA interactome of nitrogen starved *Escherichia coli* uncovers a conserved post-transcriptional regulatory axis required for optimal growth recovery

**DOI:** 10.1101/2023.10.02.560498

**Authors:** Josh McQuail, Gianluca Matera, Tom Gräfenhan, Thorsten Bischler, Per Haberkant, Frank Stein, Jörg Vogel, Sivaramesh Wigneshweraraj

## Abstract

The RNA binding protein Hfq has a central role in the post-transcription control of gene expression in many bacteria. Numerous studies have mapped the transcriptome-wide Hfq-mediated RNA-RNA interactions in growing bacteria or bacteria that have entered short-term growth-arrest. To what extent post-transcriptional regulation underpins gene expression in growth-arrested bacteria remains unknown. Here, we used nitrogen (N) starvation as a model to study the Hfq-mediated RNA interactome as *Escherichia coli* enter, experience, and exit long-term growth arrest. We observe that the Hfq-mediated RNA interactome undergoes extensive changes during N starvation, with the conserved SdsR sRNA making the most interactions with different mRNA targets exclusively in long-term N-starved *E. coli*. Taking a proteomics approach, we reveal that in growth-arrested cells SdsR influences gene expression far beyond its direct mRNA targets. We demonstrate that the absence of SdsR significantly compromises the ability of the mutant bacteria to recover growth competitively from the long-term N-starved state and uncover a conserved post-transcriptional regulatory axis which underpins this process.

## INTRODUCTION

In all three domains of life, post-transcriptional regulation of mRNA by small regulatory RNA (sRNA) represents an important mechanism to fine tune the flow of genetic information. In gram-negative bacteria such as *Escherichia coli*, sRNAs post-transcriptionally regulate more than half of all mRNAs, and many of these interactions are facilitated by the activity of an RNA binding protein (RBP), especially Hfq and ProQ (1-4).

Hfq is a hexameric Sm/Lsm-type RBP that is conserved in many bacteria. Both the protein itself and its associated sRNAs have been studied in most detail (1,5). Most Hfq-dependent sRNAs act at the ribosome binding site (RBS) by short imperfect base pairing, most commonly to repress but in some cases also to activate mRNA translation. (6-9) Further, sRNA can bind internal to a target mRNA, or within its 3’UTR. This type of interaction can either protect a ribonuclease cleavage site and thereby stabilises the mRNA to maximise chances of it being translated (10,11) or contribute to the recruitment of the mRNA to the RNA degradosome complex for degradation (12).

Individual sRNAs typically regulate multiple target mRNAs and, conversely, a target mRNA can be regulated by multiple different sRNAs. Further, some sRNAs interact with other sRNAs; this process, called ‘sponging’, represents a mechanism of regulating sRNAs by sequestering the sRNA in an inactive complex or causing their degradation (13). Additionally, sRNAs can be sponged by mRNAs (13), preventing the regulatory action of the sRNA, whilst having little regulatory effect on the mRNA. Together, this results in a complex network usually referred to as the Hfq-mediated RNA-RNA interactome, which if captured by a suitable method will predict a large part of the post-transcriptional regulation of gene expression taking place under the growth condition of interest. Recently, several methods have been developed for the capture of RNA-RNA interactomes and successfully applied to *Escherichia coli*, *Salmonella enterica*, *Vibrio cholerae*, *Staphylococcus aureus*, *Clostridioides difficile* and *Pseudomonas aeruginosa* (14-22). Importantly, these studies have underscored the view that post-transcriptional regulation confers enormous regulatory flexibility, fine-tuning and diversity to bacterial gene expression. One of these methods, RIL-seq (RNA interaction by ligation and sequencing) is a recently developed methodology, which allows genome wide mapping of RNA-RNA interactions through ligating interacting RNA molecules on RNA binding proteins (RBPs) such as Hfq (23).

Our current understanding of Hfq-mediated RNA-RNA interactomes is restricted to actively growing bacteria, bacteria experiencing iron limitation or bacteria that have entered short-term growth-arrest (4,18,21). It is widely accepted that active growth is a luxury that bacteria can only occasionally afford. Whether in a host during infection or any other natural environment, bacteria frequently exist in a growth-arrested state. To what extent Hfq-mediated RNA interactomes contribute to gene expression in growth-arrested bacteria or recovery remains unknown. Nitrogen (N) is an essential constituent for the biosynthesis of proteins and nucleic acids and N-starved bacteria are prevalent in many mammalian-intestinal systems and several fresh water-, terrestrial- and marine-ecosystems (24,25). Whereas the transcriptional basis of the adaptive response to N starvation has been broadly studied (26,27), its post-transcriptional regulatory basis is relatively poorly understood, and only one sRNA is linked the adaptive response to N starvation (see later). To address the gaps in our understanding of the roles of the Hfq-mediated RNA-RNA interactome in growth-arrested bacteria and in the adaptive response to N starvation, we applied RIL-seq to identify and compare the Hfq-mediated RNA-RNA interactome in *E. coli* at four different growth states: bacteria growing exponentially in N replete conditions (N+), short- (∼20 min; N-) and long- (∼24 h; N-24) term N-starved bacteria and bacteria ∼2 h following growth-recovery from N starvation (N-24+2). We reveal that the Hfq mediated RNA-RNA interactome is extensive and temporally dynamic in growth-arrested bacteria. By comparing how the Hfq mediated RNA-RNA interactome impacts gene expression at a proteome wide scale in long-term growth-arrested bacteria, we uncover a conserved post-transcriptional regulatory axis in growth-arrested bacteria that underpins growth recovery from long-term N starvation.

## MATERIALS AND METHODS

### Bacterial strains and plasmids

The Δ*sdsR E. coli* strain MG1655 was constructed using λ red recombination. The Δ*sdsR* and Δ*sdsR*Δ*yeaG* BW25311 mutant *E. coli* strains were created by P1 phage transduction of the Δ*sdsR* deletion into either WT or Δ*yeaG* BW25311 strains. All plasmids used in this study are listed in Table S1. p5UTR-*yeaG,* containing a translational GFP fusion of the regulatory region of *yeaG* was constructed as described previously (28) using PCR products amplified from gDNA (positions -93 to +63 relative to AUG; A is +1). Inserts were restricted with NheI and NsiI and ligated into an equally treated pXG10 plasmid backbone. Plasmid variant p5UTR-*yeaG*^MUT^ was constructed by PCR-based site-directed mutagenesis of p5UTR-*yeaG*. pSdsR was constructed using pKF68-3 (29) as a template, with the *Salmonella sdsR* changed for *E. coli sdsR* using PCR products amplified from gDNA and constructed by Gibson assembly (30). pBR322-*sdsR* was constructed by Gibson assembly using PCR product amplified from the region surrounding and including *sdsR* (∼350 bp upstream to ∼250 bp downstream of *sdsR)*.

### Bacterial growth conditions

Bacteria were grown in Gutnick minimal medium (33.8 mM KH_2_PO_4_, 77.5 mM K_2_HPO_4_, 5.74 mM K_2_SO_4_, 0.41 mM MgSO_4_) supplemented with Ho-LE trace elements (31), 0.4% (w/v) glucose as the sole carbon source and NH_4_Cl as the sole N source. Overnight cultures were grown at 37°C, 180 rpm in Gutnick minimal medium containing 10 mM NH_4_Cl. For the N starvation experiments and recovery experiments, 3 mM NH_4_Cl was used. Unless stated otherwise, for all growth assays bacteria were subcultured in growth media in a 48-well plate for a starting OD_600nm_ of 0.05, and OD_600nm_ was measured every 15 min in either a SPECTROstar OMEGA or SPECTROstar Nano plate reader (BMG LABTECH). The proportion of viable cells in the bacterial population was determined by measuring colony forming units (CFU) from serial dilutions on lysogeny broth agar plates. Overexpression experiments used pBAD18-*yeaG*, pBAD18-*yeaG*-K426A (or pBAD18-*empty* as the empty vector control), in Gutnick minimal medium supplemented with 0.125% (w/v) L-arabinose at an OD_600nm_ of approximately 0.8, for induction of *yeaG* expression.

### RIL-seq experimental procedure

RIL-seq experiments were performed as described in (18,23). Briefly, *E. coli* strains carrying either WT or 3XFLAG tagged *hfq* were grown in Gutnick minimal medium with 3 mM NH_4_Cl until OD_600_ of 0.3 (N+), ∼20 min following growth arrest (N-), 24 h following growth arrest (N-24) and ∼2 h following addition of 10 mM NH_4_Cl to N-24 cultures (N-24+2). Approximately 80 ODs were taken from three biological replicates. The samples were cross-linked under a 256 nm UV light source and pelleted in ice-cold 1xPBS. Pellets were lysed in NP-T buffer (50 mM NaH_2_PO_4_, 300 mM NaCl, 0.05% Tween, pH 8.0) supplemented with 1:200 protease inhibitor cocktail Set III, EDTA-free (Calbiochem, #539134) and RNase inhibitor (final concentration of 0.1 U/μL) (Takara, #2313A). Lysates were incubated with anti-Flag (M2 monoclonal antibody, Sigma-Aldrich, #F1804) bound protein A/G magnetic beads (Thermo-Fisher) for 2 h at 4°C with rotation followed by three washing steps with lysis buffer. Beads were treated with an RNase A/T1 mix for 5 min at 22°C in an RNase-inhibitor free lysis buffer. Samples were washed three times with lysis buffer supplemented with 3.25 μL of SUPERase In RNase inhibitor (Thermo-Fisher, #AM2696, 20 U/μL). The trimmed ends of RNAs were cured by PNK treatment (New England Biolabs, #M0201) for 2 h at 22°C with agitation, followed by 2 washing steps at 4°C. Hfq-bound RNA duplexes were ligated with T4 RNA ligase I enzyme in the following buffer: 8 μL T4 ligase buffer, 7.2 μL DMSO, 0.8 μL ATP (100 mM), 32 μL PEG 8000, 1.2 μL RNase inhibitor, 23.6 μL of water, 140 U of T4 RNA ligase I enzyme (New England Biolabs, #M0437M). Samples were incubated O/N at 22°C with agitation, followed by three steps of washing with lysis buffer, at 4°C. The RNAs were eluted from beads with a proteinase K (Thermo Fisher Scientific,#2313A) digestion for 2 h at 55°C followed by LS <u>Trizol</u> extraction, as per manufacturer instruction. Purified RNA was resuspended in 7 μL of nuclease-free water and quality controlled on a Bioanalyzer picoRNA chip. RNA samples were sent to Vertis Biotechnologie AG for downstream processing. Briefly, cDNA synthesis was conducted with oligonucleotide (dT)-adapter primers and M-MLV reverse transcriptase, followed by PCR amplification using TruSeq-designed primers from Illumina guidelines. cDNA was sequenced with an Illumina NextSeq 500.

### RIL-seq analysis and data presentation

Sequenced fragments were mapped to *E. coli* str. K-12 substr. MG1655 (U00096.3). Data analysis was performed as described by (18,23) using the RIL-seq software (https://github.com/asafpr/RILseq). Summary statistics for sequencing and mapping can be found in Table S2. Coding regions were termed CDS, annotation of 5′UTRs and 3′UTRs (termed 5UTR and 3UTR in figures and tables) was based on annotation in EcoCyc. In cases where the transcription start or termination sites of a gene were unknown (termed EST3UTR/EST5UTR for estimated UTRs), the UTRs were considered as the regions 100 nt upstream the ATG and downstream the stop codon (or shorter if these regions spanned another transcript or were more likely to be a UTR of the neighbouring transcript). Intergenic regions were termed IGR if their boundary genes were not in the same transcript or IGT if the two genes were part of the same transcript. Small RNAs that are candidate regulatory RNAs as denoted by EcoCyc were grouped as sRNAs. Transcripts antisense to genes or to IGT were termed AS and AS_IGT respectively. The same annotation was maintained in all tables and figures unless noted otherwise. ygaM.EST3UTR was re-annotated as an sRNA. For the purpose of certain data analysis and data interpretation glnA.3UTR was considered to be GlnZ, ybaP.EST3UTR was considered to be ChiX, glmZ.hemY was considered to be GlmZ and nlpD was generally considered to be rpoS. Interaction types were defined as follows: mRNA:sRNA for chimeras with one sRNA and either a CDS, 3UTR, 5UTR, EST3UTR, EST5UTR or IGT, sRNA:sRNA for chimeras with two sRNA, sRNA:tRNA for chimeras with an sRNA and a tRNA, and other interactions were defined as all other interactions not categorised by the previous definitions (including all those involving AS or IGR fragments). Principle component analysis was performed on total RIL-seq datasets following both normalisation and standardisation of raw chimera counts using the factoextra package in R 4.2.3 and plotted using the scatterplot3d package in R 4.2.3. For further downstream analysis and interpretation only interactions that were detected with at least 30 chimeras, and in least two replicates from the same time-point were considered. Chimera data from individual samples can be found in Table S3, and summary data from all samples can be found in Table S4. For certain comparisons between time-points and/or replicates the number of chimeras for each interaction were normalised such that the total number of chimeras in each replicate and time-point were the same (130677). Circos plots were generated using shinyCricos (https://venyao.xyz/shinyCircos-V1/) and then edited further in Adobe Illustrator 26.5. Further information on specific Cricos plots are provided in their respective figure legends. sRNA-target binding predictions were performed using IntraRNA (32). Coverage plots of chimeras were generated using the script generate_BED_file_of_endpoints.py of the RIL-seq computational pipeline; the bed files were visualized with IGV 2.12.2, and further processed with Adobe Illustrator 26.5.

### Proteomics sample preparation

The proteomics analysis presented in the worked was performed as part of a larger proteomics experiment consisting for WT and 4 different mutant *E. coli* strains. For total-proteome analysis WT and Δ*sdsR E. coli* were grown to N-24, and 20 ml of culture was washed twice with ice cold PBS and flash frozen in liquid nitrogen before further processing. Pellets were suspended in 500 μl of lysis buffer (100 mM Tris-HCl (pH7.9), 150 mM NaCl, 1.5% SDS, 2X cOmplete Protease Inhibitor mix (Roche)) and sonicated on ice for a total of 10 min at 30% intensity, in 10 s intervals. Insoluble protein and cell debris was pelleted by centrifugation for 20 min and supernatant was collected. All samples were adjusted to an equal protein concentration with more lysis buffer. 10 µg of lysate derived from our control (WT strain), Δ*sdsR* strain, and other samples not discussed in this study, were subjected to an in-solution tryptic digest using a modified version of the Single-Pot Solid-Phase-enhanced Sample Preparation (SP3) protocol (33,34). 1% SDS-containing lysates were added to Sera-Mag Beads (Thermo Scientific, #4515-2105-050250, 6515-2105-050250) in 10 µl 1 5% formic acid and 30 µl of ethanol. Binding of proteins was achieved by shaking for 15 min at room temperature. SDS was removed by 4 subsequent washes with 200 µl of 70 % ethanol. Proteins were digested overnight at room temperature with 0.4 µg of sequencing grade modified trypsin (Promega, #V5111) in 40 µl Hepes/NaOH, pH 8.4 in the presence of 1.25 mM TCEP and 5 mM chloroacetamide (Sigma-Aldrich, #C0267). Beads were separated, washed with 10 µl of an aqueous solution of 2% DMSO and the combined eluates were dried down. Peptides were reconstituted in 10 µl of H2O and reacted for 1 h at room temperature with TMT16pro labelling reagent (Thermo Scientific, #A44522). To this end, 50 µg of TMT16pro label reagent were dissolved in 4 µl of acetonitrile and added to the peptides. Excess TMT reagent was quenched by the addition of 4 µl of an aqueous 5% hydroxylamine solution (Sigma, 438227). Peptides were reconstituted in 0.1% formic acid and equal volumes were mixed. Mixed peptides were purified by a reverse phase clean-up step (OASIS HLB 96-well µElution Plate, Waters #186001828BA). Peptides were subjected to an off-line fractionation under high pH conditions (33) yielding 6 fractions.

### LC-MS/MS analysis

Peptides were separated using an Ultimate 3000 nano RSLC system (Dionex) equipped with a trapping cartridge (Precolumn C18 PepMap100, 5 mm, 300 μm i.d., 5 μm, 100 Å) and an analytical column (Acclaim PepMap 100. 75 × 50 cm C18, 3 mm, 100 Å) connected to a nanospray-Flex ion source. The peptides were loaded onto the trap column at 30 µl per min using solvent A (0.1% formic acid) and eluted using a gradient from 2 to 80% Solvent B (0.1% formic acid in acetonitrile) over 90 min at 0.3 µl per min (all solvents were of LC-MS grade). The Orbitrap Fusion Lumos was operated in positive ion mode with a spray voltage of 2.4 kV and capillary temperature of 275°C. Full scan MS spectra with a mass range of 375–1500 m/z were acquired in profile mode using a resolution of 120,000 with a maximum injection time of 50 ms, AGC operated in standard mode and a RF lens setting of 30%. Fragmentation was triggered for 3 s cycle time for peptide like features with charge states of 2–7 on the MS scan (data-dependent acquisition). Precursors were isolated using the quadrupole with a window of 0.7 m/z and fragmented with a normalized collision energy of 34%. Fragment mass spectra were acquired in profile mode and a resolution of 30,000 in profile mode. Maximum injection time was set to 94 ms or an AGC target of 200%. The dynamic exclusion was set to 60 s.

### Proteomics data analysis

Acquired data were analyzed using FragPipe (35) and a Uniprot *E. coli* FASTA database (UP000000625, *E. coli* strain K12, ID83333, 4402 entries, last modified October 27th 2022, downloaded January 11th 2023) including common contaminants. The following modifications were considered: Carbamidomethyl (C, fixed), TMT16plex (K, fixed), Acetyl (N-term, variable), Oxidation (M, variable) and TMT16plex (N-term, variable). The mass error tolerance for full scan MS spectra was set to 10 ppm and for MS/MS spectra to 0.02 Da. A maximum of 2 missed cleavages were allowed. A minimum of 2 unique peptides with a peptide length of at least seven amino acids and a false discovery rate below 0.01 were required on the peptide and protein level (36).

### Proteomics statistical data analysis

The raw output files of FragPipe (protein.tsv – files, (35)) were processed using the R programming language (ISBN 3-900051-07-0). Contaminants were filtered out and only proteins that were quantified with at least two unique peptides were considered for the analysis. 2156 proteins passed the quality control filters. Log2 transformed raw TMT reporter ion intensities were first cleaned for batch effects using the ‘removeBatchEffects’ function of the limma package (37) and further normalized using the vsn package (variance stabilization normalization - (38)). Proteins were tested for differential expression using the limma package. The replicate information was added as a factor in the design matrix given as an argument to the ‘lmFit’ function of limma. Limma analysis for Δ*sdsR* as compared to WT can be found in Table S5.

### Immunoblotting

Immunoblotting was conducted in accordance to standard laboratory protocols, with primary antibodies incubated overnight at 4°C. A rabbit anti-YeaG antibody was produced previously (39) and used at a dilution of 1:2,000. The following other antibodies were used: mouse monoclonal anti-RpoA (Biolegend, WP003) at 1:1,000 dilution, HRP Goat anti-mouse IgG (BioLegend, 405306) at 1: 10, 000 dilution and HRP Goat anti-rabbit IgG (GE Healthcare, NA934) at 1:10,000 dilution. ECL Prime Western blotting detection reagent (GE Healthcare, RPN2232) was used to develop the blots, which were analysed on the ChemiDoc MP imaging system and bands quantified using Image Lab software.

### SdsR target binding site analysis

Binding site designations for mRNA targets of SdsR identified in the RIL-seq dataset were determined based on the peak of chimeric alignments to their target RNA in IGV. Designations were defined as follows: RBS – 40 bp upstream to 15 bp downstream of the AUG; Internal + 3’UTR – any site more than 15 base pairs following the AUG and before any known terminator (this designation includes targets with multiple internal binding sites); 5’UTR - in the 5’UTR but more than 40 bp upstream of the AUG; multiple sites - when chimeras map to more than one designation; poorly defined - when chimeras mapped to a very broad region of the mRNA or were found in an intergenic region. Targets that mapped in an intergenic region between transcripts were excluded from binding site analysis.

### Scanlag

Bacteria were grown in 3 mM Gutnick minimal medium to N-24, washed twice in 1 ml PBS, diluted between 10^−5^ to 10^−6^, and 100 μL spread on Gutnick minimal media containing 0.4% glucose, 3 mM NH_4_Cl, Ho-LE trace elements and 1.5% (wt/vol) agar. Plates were incubated at 33°C in a standard office scanner (Epson Perfection V370 photo scanner, J232D) placed in an incubator, and images were taken every 20 min over a 48 h period. Analysis of appearance time and apparent growth rate of colonies was adapted from Levin-Reisman et al. (40) using a modified code available at https://github.com/mountainpenguin/NQBMatlab.

### Competition assay

For bacterial competition assays, bacteria were grown to the desired growth state (N- or N-24) in Gutnick minimal media supplemented with 0.4% glucose and 3 mM NH_4_Cl. In all competition assays the Δ*sdsR* strain was resistant to kanamycin, and the WT was not. For competition during recovery growth, WT and Δ*sdsR* bacteria were both inoculated at a starting OD_600mn_ of 0.025 in Gutnick minimal media supplemented with 0.4% glucose and 3 mM NH_4_Cl, for a total starting OD_600nm_ of 0.05. CFUs were measured by serial dilution, at the point of inoculation and at an OD_600nm_ of 0.4 and 0.8 - plating on both LB agar and LB agar with 50 μg/ml of kanamycin; the CFUs on the plates containing kanamycin gives the number of Δ*sdsR* bacteria, and the CFU on the plates with kanamycin give the total number of bacteria - the number of WT bacteria can then be calculated by subtracting the number of Δ*sdsR* bacteria from the total number of bacteria. For competition during stationary phase, both WT and Δ*sdsR* bacteria were resuspended in Gutnick minimal media supplemented with 0.1% glucose and no nitrogen source; OD_600nm_ was measured and wild-type and Δ*sdsR* bacteria were combined to an equal OD_600nm_ for a total final volume of 20 ml. CFUs for total and Δ*sdsR* bacteria were then determined at regular intervals as above.

### SdsR YeaG-GFP reporter assay

sRNA-target regulation assays were performed as previous, with some modifications (28). Plasmids pSdsR and pCONT contained either SdsR or a ∼50 nt nonsense RNA fragment from an IPTG inducible promoter. Plasmids p5UTR*-yeaG* contained the 5’UTR and translational start codon of *yeaG* (positions -93 to +63 relative to AUG; A is +1) N-terminally fused to GFP from a constitutively active promoter, and p5UTR*-yeaG^MUT^* contained the same with the CG at residues -28/27 relative to the AUG mutated to GC. The required combinations of plasmids were transformed into Δ*sdsR* MG1655 *E. coli* and overnight cultures were grown in LB containing 50 μg/ml kanamycin, 100 μg/ml ampicillin and supplemented with 0.4% glucose to repress expression of SdsR from pSdsR. All bacterial strains were then inoculated into 0.2 ml LB containing 50 μg/ml kanamycin, 100 μg/ml ampicillin, either with or without 0.1 mM IPTG for induction of SdsR expression, at a starting OD600nm of 0.05, in a black-walled 96-well plate. Cultures were then grown in a SPECTROstar OMEGA plate reader at 37°C for 10 h. OD_600nm_ and GFP fluorescence were then measured.

### Lag time and doubling time

Lag time was calculated by generating a straight-line graph of the exponential growth section of each growth curve, and was defined as the intersection of the straight-line graph with the x axis. The doubling time was determined from the slope of logarithmic growth function.

### RNA-sequencing and analysis

WT and Δ*sdsR* bacteria were harvested at N-24. RNA was extracted using the RNAsnap protocol (41). Three biological replicates of each strain were taken and mixed with a phenol:ethanol (1:19) solution at a ratio of 9:1 (culture:solution) before harvesting the bacteria immediately by centrifugation. Pellets were resuspended in RNA extraction solution (18 mM EDTA, 0.025% SDS, 1% 2-mercaptoethanol, 95% formamide) and lysed at 95°C for 10 min. Cell debris was pelleted by centrifuged. RNA was precipitated by addition of 1/10 volume of 3 M sodium acetate (pH 5.2) and 3 volumes of 100% ethanol, followed by incubation at -80°C for at least 1 h. RNA was washed with 75% ethanol and resuspended in water. Extracted RNA was sent to the Core Unit Systems Medicine at the University of Würzburg for next-generation sequencing and alignment to the reference *E. coli* K12 MG1655 (U00096.3). Total read counts were used to normalize the data, the number of reads mapping to each gene was calculated, and a matrix of read counts was generated. The matrix was analysed using the DESeq2 BioConductor package for differential gene expression analysis (Table S6) (42).

## RESULTS

### RIL-seq reveals an extensive and dynamic RNA-RNA interactome in N starved *E. coli*

RIL-seq allows isolation and sequencing of ligated RNA-RNA pairs through immunoprecipitation of Hfq-RNA complexes (see Materials and Methods). To immunoprecipitate Hfq, we used an *E. coli* strain, which contained a 3xFLAG-tag sequence fused C-terminally to *hfq* at its normal chromosomal location. Control experiments revealed that the growth dynamics (Figure S1A) and survival (Figure S1B) during growth arrest of *E. coli* strains without and with 3xFLAG were indistinguishable, indicating that the modification to Hfq did not compromise its activity under our experimental conditions. We then performed RIL-seq in N+, N-, N-24 and N-24+2 *E. coli*, grown in a defined minimal growth medium with a limiting (∼3 mM) amount of ammonium chloride as the only N source. N+ bacteria were sampled from the exponential phase of growth at an OD_600_ of ∼0.3 (Figure 1A). Growth arrest occurs as soon as the ammonium chloride in the media runs outs (27,43-45) and we considered bacteria ∼20 min after N run-out to be in the N-state. The N-24 bacteria were sampled ∼24 h following N run-out. Growth recovery was induced by supplying ∼3 mM ammonium chloride to N-24 bacteria and N-24+2 bacteria were collected following 2 h. Details on the sequencing depth and mapping statistics are listed in Table S2. The data was analysed as described in by Matera et al., (16) (Table S3 & S4). Statistically significant chimeras from the Hfq precipitated showed an abundance of sRNAs and mRNAs over the control sample (Figure 1B and Figure S1C). Principal component analysis (PCA) showed that Hfq-mediated RNA-RNA interactomes from the individual replicates from each growth stage clustered together, underscoring the stringency and reproducibility of the sampling and experimental approach (Figure 1C). We note that chimeras from N+ and N-24+2 bacteria were closely clustered, contrasting the chimeras from N- and N-24 bacteria, which were represented by two distinct clusters, suggesting that the Hfq-mediated RNA-RNA interactome dramatically differ between growing and growth-arrested bacteria and in bacteria in different stages of growth-arrest (see later). Only RNA pairs represented by at least 30 chimeric fragments in at least two replicates were considered for downstream analysis, giving ∼315, ∼354, ∼379, ∼279 interactions in N+, N-, N-24 and N-24+2 bacteria, respectively (Table S2 & S3). Most of these corresponded to sRNA-mRNA chimeras (mRNA collectively defined here as coding sequences [CDSs], 5′ UTRs, 3’UTRs and intergenic regions within transcripts [IGTs]) (Figure S1D). As with previous Hfq-mediated RNA-RNA interactomes (see Introduction), we also detected numerous interactions between two sRNAs (i.e., sponging interactions), consistent with the idea that Hfq-mediated post-transcriptional regulation extends far beyond its canonical role in mRNA regulation (13,46) (Figure 1D and Figure S2).

**Figure 1.**
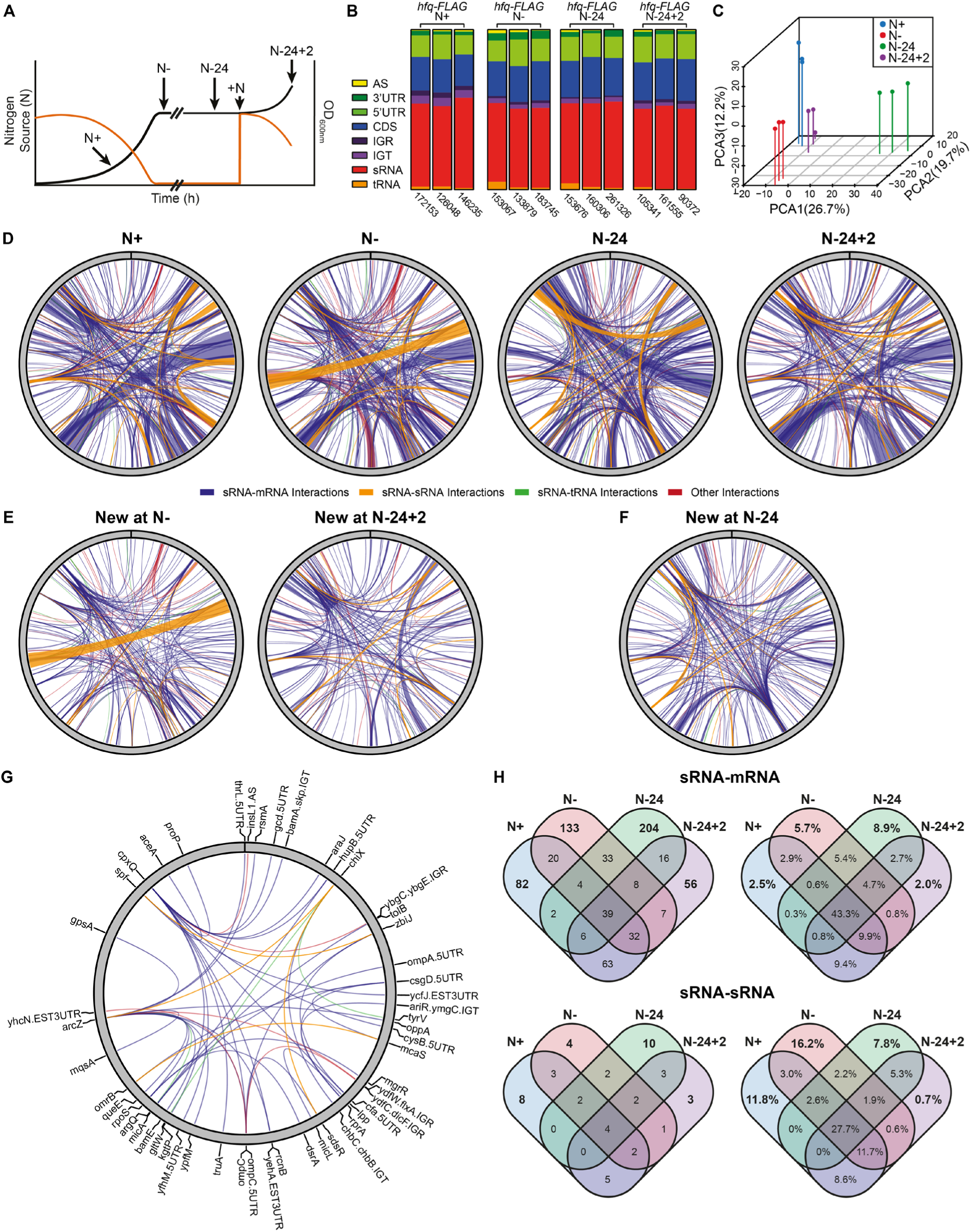
RIL-seq reveals an extensive and dynamic Hfq-mediated RNA interactome in N starved *E. coli*. **(A)** Schematic representation of experimental design, indicating growth states used throughout this study. **(B)** Relative frequencies of each RNA type found in chimeric fragments, in individual replicates across all time points for *hfq-FLAG* datasets. **(C)** Principal component analysis of complete chimeric fragment datasets from each replicate. **(D)** Circos plots of RIL-seq interactions that are represented by at least 30 chimeric fragments in two individual replicates, mapped to the *E. coli* K12 MG1655 genome. The thickness of each connection is proportional to the average number of chimeras detected for a given interaction across the three replicates. sRNA:mRNA, sRNA:sRNA, sRNA:tRNA and other interactions are represented by blue, orange, green and red lines respectively. **(E)** Circos plots of the RIL-seq interactions detected at N-but not N+ (left), and N-24+2 but not N-24 (24). **(F)** Circos plot of the RIL-seq interactions detected at N-24 but not N-. **(G)** Circos plot of RIL-seq interactions detected at all time points. Thickness of connections are not weighted by the number of chimeras. The RNA involved in each connection are shown. **(H)** Venn diagrams showing the condition specificity of sRNA:mRNA and sRNA:sRNA interactions. Diagrams on the left show the number of interactions, diagrams on the right are weighted by the number of detected chimeras for each interaction, shown as a percentage of the entire dataset.

Circos plot representations of the sRNA-mRNA, sRNA-sRNA, sRNA-tRNA and other (e.g., sRNA interacting with intergenic region between two transcripts or antisense RNA) showed that the Hfq-mediated RNA-RNA interactome is dominated by sRNA-mRNA and sRNA-sRNA interactions across all growth states (Figure 1D, Figure S2). As expected, the Hfq-mediated RNA interactomes substantially changed upon transition from N+ to N- (225 new at chimeras at N-) and N-24 to N-24+2 (187 new chimeras at N-24+2) growth states (Figure 1E). Notably, the Hfq-mediated RNA-RNA interactome also substantially differed in bacteria between the two different growth-arrested states, i.e., between N- and N-24 states (∼266 new chimeras at N-24; Figure 1F). Conversely, we noted that several sRNA-mRNA (∼39), sRNA-sRNA (∼4), sRNA-tRNA (∼3) and other (∼5), interactions were present across all four growth states, which we termed the ‘core’ Hfq-mediated RNA interactome in our condition (Figure 1G). However, the abundance of some of the core RNA-RNA interactions differed at each growth state (Figure S1E). The number of Hfq-mediated sRNA-mRNA and sRNA-sRNA interactions that are unique to or common between each growth state is summarised in Figure 1G. Overall, we conclude that the Hfq-mediated RNA interactome is extensive and highly dynamic in *E. coli* experiencing N starvation.

### An elaborate post-transcriptional regulatory basis of the adaptive response to N-starvation in *E. coli*

*E. coli* cells adapt to N starvation by activating the nitrogen regulation (47) stress response, resulting in the expression of 20 operons (26,27). The master transcription regulator of the Ntr response is NtrC of the NtrBC two-component system. The activation of the Ntr response results in a large scale reprogramming of gene expression at the transcriptional level because NtrC activated genes include Nac, a global transcription factor affecting the transcription of ∼80-100 genes (48) and RelA, the enzyme responsible for the synthesis of guanosine tetraphosphate (ppGpp), which acts directly on the RNA polymerase and alters its activity and promoter specificity (49). In contrast, the extent to which post-transcriptional regulation contributes to the adaptive response to N starvation is not fully understood. Recently, the sRNA GlnZ was shown to promote cell survival by regulating genes linked to N and carbon flux in N starved *E. coli* in an Hfq-dependent manner (50,51). In MS2-affinity purification and RNA sequencing (MAPS) experiments using GlnZ as bait, conducted in short-term N-starved *E. coli* (i.e., a condition comparable to N-), GlnZ interacted with numerous mRNA targets (51). In Figure S3A, we show the GlnZ regulon obtained by RIL-seq in *E. coli* bacteria in N+, N-, N-24 and N-24+2 states. The mRNAs of the 11 genes, which were enriched in the MAPS experiments are shown in green. Interestingly, RIL-seq analysis revealed 5 new potential GlnZ targets, which also included an interaction with the *nac* mRNA at N-. This suggests that the NtrC and Nac regulon are potentially also coupled at the post-transcriptional level (see Discussion). Further, we interrogated the RIL-seq data from all four growth states for sRNAs interacting with mRNAs of genes that belong to the Ntr regulon (i.e., genes directly activated by NtrC). The results in Figure S3B reveal that the expression of a total of 13 of the 20 NtrC dependent operons (shown in blue) are potentially affected by 16 different sRNAs (shown in red) in N+ (3 operons), N- (12 operons) and N-24 states (4 operons) and N-24+2 (2 operons) bacteria. Overall, the results highlight that post-transcriptional regulation of gene expression is likely to be an important, yet unexplored, facet of the adaptive response to N starvation in *E. coli*.

### The conserved sRNA SdsR has a substantial post-transcriptional regulatory influence in *E. coli* experiencing long-term N starvation

To understand post-transcriptional control mechanisms that underpin gene expression in long-term growth arrested bacteria, we focused on the Hfq-mediated RNA-RNA interactome in N-24 *E. coli*. Figure 2A shows the top 10 sRNAs that contribute to the Hfq-mediated RNA-RNA interactome across all four growth states, with the SdsR sRNAs contributing to almost a third (∼31.5%) of chimeras. SdsR, which is ∼100 nucleotide long, is one of the most highly conserved enterobacterial sRNAs (Figure S4A). Its transcription is dependent on the RNAP containing the general stress response promoter specificity factor RpoS and as such SdsR is produced when cells encounter stress and/or growth-arrest (29,52). In previous RIL-seq experiments, the Hfq-mediated interactome of SdsR consisted of ∼92 interactions in *E. coli* from stationary phase of growth in lysogeny broth (i.e., a growth state comparable to N-) and only one Hfq-mediated RNA-RNA interaction involving SdsR was detected in *E. coli* from the exponential phase of growth (i.e., comparable to the N+ bacteria used here) (18). Consistent with previous results, only 4, 31 and 6 chimeras involving SdsR (representing ∼0.3, ∼4.9 and ∼0.9 % of the Hfq-mediated RNA interactome) were detected in N+, N- and N-24+2 bacteria, respectively. In N-24 bacteria, SdsR interacted with 124 mRNA targets (Figure 2B). To establish how many of these interactions were productive, we compared the total proteomes of wild-type and Δ*sdsR* bacteria at N-24 (Figure 2C). Neither the growth dynamics of wild-type and Δ*sdsR* (Figure S5A) nor the proportion of viable cells in the N-24 population of wild-type and Δ*sdsR* (Figure S5B) were substantially different. We identified ∼50% (2,156/4,328) of total *E. coli* proteins in the proteomes of wild-type and Δ*sdsR* bacteria. Notably, the absence of SdsR resulted in the differential expression of ∼70% (p-value ≤0.05) of the identified proteins in Δ*sdsR* compared to wild-type bacteria. Of the 124 potential mRNA targets of SdsR, the proteins of 90 were identified in the proteomics data, of which 69 were designated as significantly differentially expressed in the Δ*sdsR* proteome (shown red in Figure 2C). Of these 69 proteins, SdsR bound to 25 of the corresponding mRNAs at or overlapping the RBS (defined here as 40 bp upstream to 15 bp downstream of the AUG codon), 35 mRNAs internally or in the 3’ UTR, 2 mRNAs in the 5’UTR not overlapping the RBS and 6 mRNAs to multiple binding sites within the mRNA, suggesting that SdsR can influence the post-transcriptional fate of its mRNA targets in several different ways (see Introduction) in long-term N starved *E. coli*.

**Figure 2.**
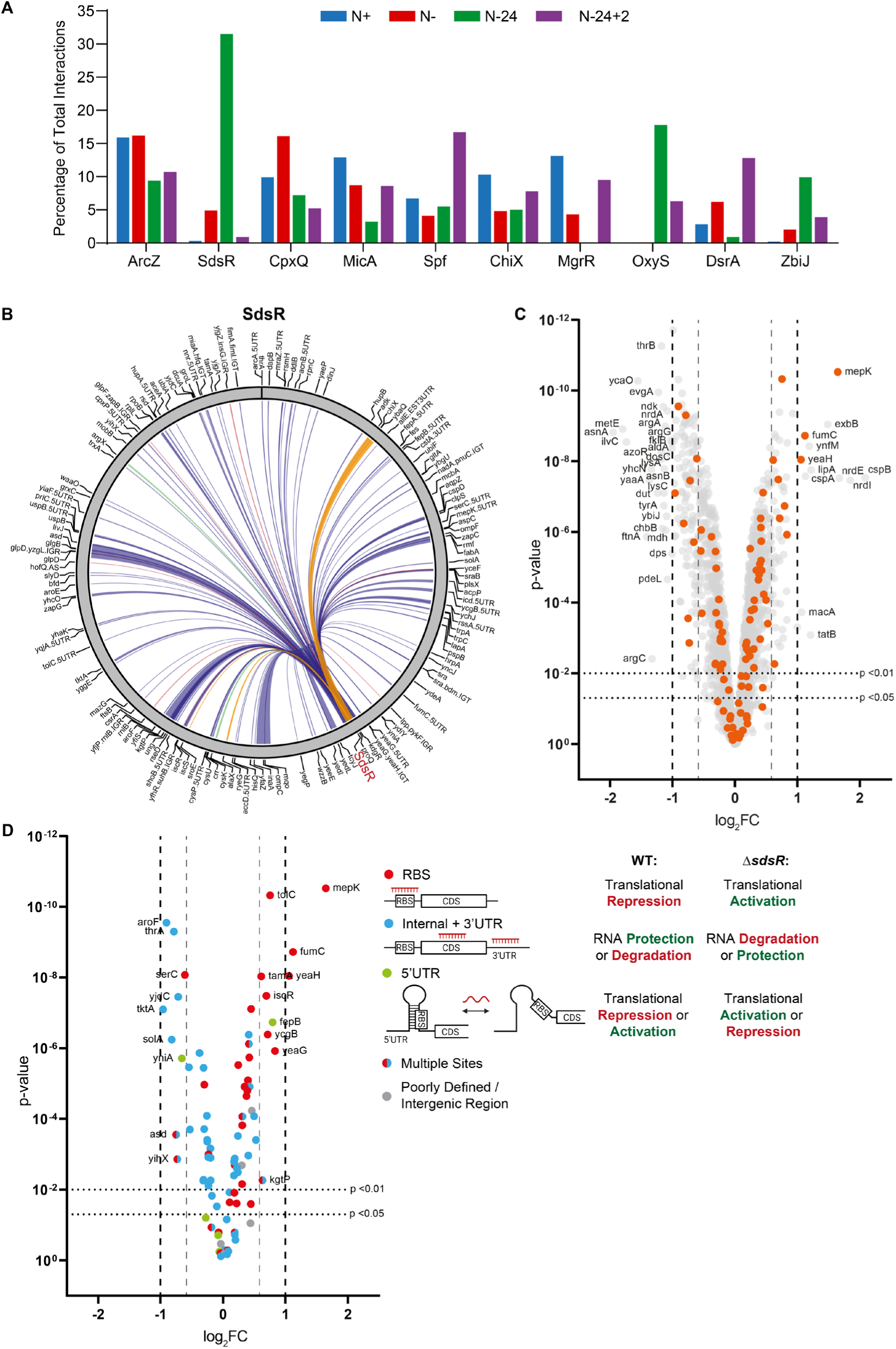
The conserved sRNA SdsR has a substantial post-transcriptional regulatory influence in *E. coli* experiencing long-term N starvation. **(A)** Graph showing the relative abundance of the top ten most represented sRNA in RIL-seq chimeras at each time point. **(B)** Circos plot of interactions involving SdsR at N-24 that are represented by at least 30 chimeric fragments in two individual replicates. The thickness of each connection is proportional to the average number of chimeras detected for a given interaction across the three replicates. sRNA:mRNA, sRNA:sRNA, sRNA:tRNA and other interactions are represented by blue, orange, green and red lines respectively. **(C)** Volcano plot of differential protein levels in N-24 Δ*sdsR* bacteria shown as a log_2_ change from wild-type bacteria. Proteins differentially expressed more than 1 log_2_ (i.e., a greater than 2-fold change) are indicated, and those proteins whose mRNA interacted with SdsR in the RIL-seq dataset are coloured orange. **(D)** As above, showing only proteins whose mRNA interacted with SdsR, coloured by the SdsR binding site as determined though RIL-seq. Interaction regions overlapping the RBS (defined here as 40bp upstream to 15bp downstream of the AUG) are shown in green, interactions internal to the coding sequence or in the 3’ UTR are shown in red, interactions in the 5’UTR not overlapping the RBS are shown in blue, targets with multiple binding sites within the mRNA are shown with split colour and poorly defined and intergenic regions between transcripts are shown in grey. Schematic representation of the categories of sRNA binding site, and the expected regulatory outcome on protein levels in the wild-type and Δ*sdsR* bacteria are schematically shown.

We expected that proteins of mRNAs to which SdsR bound at or overlapping the RBS to be upregulated in the Δ*sdsR* proteome compared to the wild-type proteome, because sRNA interference at the RBS usually reduces translation (see Introduction). We expected that proteins of mRNAs to which SdsR bound internally or at the 3’ UTR to be either downregulated or upregulated in the Δ*sdsR* proteome compared to the wild-type proteome, because sRNA action at these sites can result in the protection of the targeted mRNA from degradation or enhanced degradation of the targeted mRNA, respectively (see Introduction). Consistent with this view, ∼88% (22/25) of the detected proteins where SdsR was bound to sites at or overlapping the RBS of their corresponding mRNAs were found to be upregulated in the Δ*sdsR* proteome (Figure 2D, red dots). However, ∼63% (22/35) and ∼37% (14/36) of the detected proteins where SdsR was bound internally or in the 3’UTR of their corresponding mRNA were found to be downregulated and upregulated, respectively, in the Δ*sdsR* proteome (Figure 2D, blue dots). Overall, we conclude that in N-24 *E. coli* (i) the regulatory influence of SdsR extends far beyond its *direct* regulatory targets and thereby *indirectly* affects diverse and broad range of cellular processes (Figure S5C), (35) a large proportion of the interactions between SdsR and its mRNA targets have a productive regulatory effect and (iii) SdsR has both a broad positive and negative regulatory effect on gene expression.

### SdsR is required for optimal growth recovery from long-term N starvation

Although SdsR affects the expression of genes associated with diverse processes in N-24 bacteria, its absence, surprisingly, has little impact on the growth or long-term survivability of Δ*sdsR* bacteria (Figure 5SA and 5SB). Therefore, we considered whether SdsR had any influence on the ability of N-24 bacteria to recover from growth-arrest. We discovered that when N-24 Δ*sdsR* bacteria were inoculated into fresh culture media, they consistently displayed an increased lag phase (here defined as the time from inoculation to the doubling of the OD_600_ reading of the culture) by ∼57 min longer than wild-type bacteria (Figure 3A, left). The increased lag phase of Δ*sdsR* bacteria could be reverted to that of wild-type bacteria when plasmid-borne SdsR was expressed from its native promoter in Δ*sdsR* bacteria (Figure 3A, inset); although we noted that harbouring the pBR322 plasmid further increases the lag phase of Δ*sdsR* bacteria by ∼45 min compared to Δ*sdsR* bacteria without any plasmid). This increased lag phase was not detected when N-Δ*sdsR* bacteria were recovered in fresh media (Figure 3A, right), demonstrating that the increased lag phase is a specific property of long-term N starved Δ*sdsR* bacteria.

**Figure 3.**
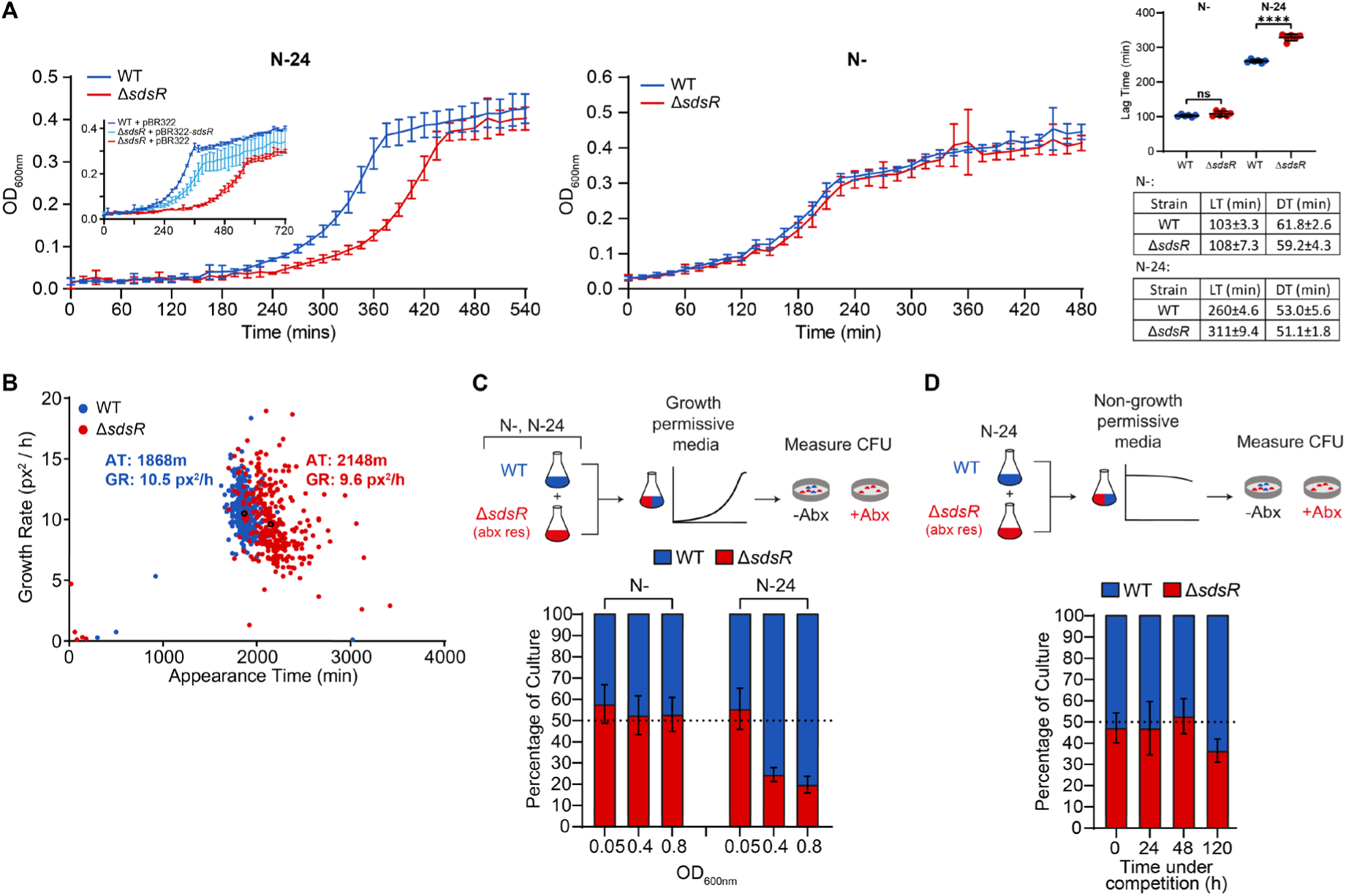
SdsR is required for optimal growth recovery from long-term N starvation. **(A)** Graphs showing recovery growth of wild-type and Δ*sdsR* bacteria following sub-culturing of 24 h (Left) and 20 min (Right) N-starved bacteria into fresh culture media. Inset shows recovery of wild-type and Δ*sdsR* bacteria containing either empty pBR322 or pBR322 expressing *sdsR* from its native promoter, following sub-culturing of N-24 bacteria into fresh culture media. Error bars represent standard deviation (n=6). Additional graph shows lag-time and tables show lag-time (LT) and doubling-time (DT). Statistical analysis performed by Welch’s T-test (****, P<0.0001). **(B)** Scanlag analysis of colony appearance time (in min) and growth rate (in pixels^2^/h (px^2^/h)) for wild-type (blue) and Δ*sdsR* (red) strains experiencing N starvation for 24 h and plated onto Gutnick agar with 3mM NH_4_Cl. Black circles represent average population growth rate and appearance time with mean values indicated. **(C)** Graph of the proportions of wild-type and Δ*sdsR* following co-inoculation of equal amounts of wild-type and Δ*sdsR* bacteria from the N- and N-24 growth states into fresh growth media. Proportions were determined by CFU measured at an OD_600nm_ of 0.05, 0.4 and 0.8 in the co-culture, following plating on LB agar with (to select only for Δ*sdsR* bacteria) and without kanamycin (to select for total number of bacteria). The experimental approach is shown schematically at the top. **(D)** As in (C), but equal proportions of N-24 wild-type and Δ*sdsR* bacteria were resuspended in Gutnick media without any NH_4_Cl, incubated for up to 120 h, with proportion of wild-type and Δ*sdsR* bacteria determined at regular intervals.

We considered whether the difference in lag phase between wild-type and Δ*sdsR* bacteria upon recovery from N-24 state was due to the presence of a subpopulation(s) of bacteria in the Δ*sdsR* population that resumed growth slowly. To investigate this, we used a method called ScanLag, which allows the measurement of the appearance time of individual colonies on a solid growth medium as a function of time following inoculation (40). We considered that if the N-24 Δ*sdsR* population consisted of one or more slow growing subpopulations compared to the wild-type population, then the appearance of individual colonies of Δ*sdsR* bacteria would occur at different times than that of wild-type bacteria. As shown in Figure 3B, when N-24 Δ*sdsR* bacteria and wild-type bacteria were plated on solid media containing ammonium chloride as the sole N source, most of the Δ*sdsR* colonies appeared at the same time, but their appearance time differed by ∼280 min compared to that of the wild-type colonies. Although, we noted that the appearance time of individual Δ*sdsR* colonies were inherently more heterogenous than that of wild-type colonies.

As bacteria exist in polymicrobial environments, we considered whether the subtle increase in lag time to growth recovery of N-24 Δ*sdsR* bacteria would lead to a competitive disadvantage when recovered in a mixed population with wild-type bacteria. To investigate this, we inoculated the same number (determined by counting colony forming units [CFU]) of Δ*sdsR* and wild-type cells from N-24 into fresh media and enumerated the CFU of Δ*sdsR* bacteria as a percentage of total bacteria in the culture. Following initial inoculation at OD_600_ ∼0.05, when Δ*sdsR* and wild-type bacteria were present at approximately equal proportions, at OD_600_ ∼0.4 and ∼0.8, the proportion of Δ*sdsR* bacteria decreased to that of ∼20% of the total population (Figure 3C). Such a difference was not detected when N-Δ*sdsR* and wild-type bacteria were co-inoculated (Figure 3C), consistent with the results shown in Figure 3A (right) when no difference in lag time between Δ*sdsR* and wild-type bacteria was observed during growth recovery. The competitive disadvantage of the Δ*sdsR* bacteria was specific to growth recovery from the N-24 state, as it was absent when N-24 Δ*sdsR* and wild-type were co-inoculated into fresh but N depleted media, which did not support growth (Figure 3D). In sum, we conclude that the post-transcriptional regulon of SdsR contributes to optimal growth recovery of long-term N-starved bacteria.

### Optimal growth recovery from long-term N starvation requires translational downregulation of the conserved *yeaGH* operon by SdsR

We interrogated the proteome of Δ*sdsR* bacteria to identify genes post-transcriptionally regulated by SdsR that might be responsible for the lag in the growth recovery of N-24 Δ*sdsR* bacteria. We focused on proteins whose translation was directly affected by SdsR and transcription of their mRNAs was activated by NtrC. We identified HisQ, YeaG and YeaH as such proteins, which were ∼1.32, ∼1.78 and ∼2.08-fold, respectively, more abundant in N-24 Δ*sdsR* bacteria than in wild-type bacteria. As we previously implicated YeaG and YeaH in the adaptive response to N starvation (44,45), we focused our analysis on these two proteins.

The products of the *yeaGH* operon are conserved in several enterobacteria (Figure S4B and Figure S4C) and the *yeaGH* operon is expressed (in addition to N starvation) when bacteria encounter diverse stresses such as low pH (53), high osmolarity (54), stationary phase (53), low sulphur (55). YeaG is a Hank’s type eukaryote-like serine/threonine kinase (56,57) but the biological function of YeaH remains unknown. In previous work, we reported that the *yeaGH* operon is activated by NtrC when *E. coli* experiences N starvation (i.e., at N-) and that YeaG and YeaH functionally cooperate and, by a yet to be characterised mechanism, contribute to quiesce metabolism in N-starved *E. coli* (44,45). Therefore, the absence of *yeaG* or *yeaH* compromises the long-term survivability of mutant bacteria under N starvation (44,45). Notably, the growth recovery phenotype of Δ*yeaG*, Δ*yeaH* and Δ*yeaGH* bacteria contrasts that of the Δ*sdsR* bacteria: The Δ*yeaG*, Δ*yeaH* and Δ*yeaGH* bacteria recover from the N-24 state with a shorter lag phase (by ∼30 min) compared to wild-type bacteria (see later and (44)).

As YeaG is experimentally more tangible than YeaH, we were previously able to purify and raise antibodies against YeaG to demonstrate that YeaG protein levels peak during first 6-9 h from the onset of N starvation but decrease by N-24 (45), suggesting a potential post-transcriptional regulatory mechanism underpinning *yeaGH* gene expression. Our new data have now revealed that *yeaGH* expression is subjected to translational downregulation by SdsR, underscoring the view that managing the intracellular levels of YeaG (and YeaH) is important for optimal growth recovery from long-term N starvation. Consistent with this view, immunoblotting of YeaG in N-24 wild-type and Δ*sdsR* bacteria revealed a substantially increased abundance of YeaG in Δ*sdsR* bacteria compared to in wild-type bacteria (Figure 4A). However, the transcription of *yeaGH* operon can be driven by two different promoters that depend on the RNAP containing the promoter specificity factor RpoS or RpoN. Inspection of the proteome of the Δ*sdsR* bacteria at N-24 reveals that RpoS and RpoN proteins are ∼34% and ∼13%, respectively, more abundant in Δ*sdsR* bacteria than in wild-type bacteria (Figure S6A). Therefore, it is possible that the increased abundance of YeaG and YeaH in Δ*sdsR* bacteria is potentially due to increased abundance of *yeaGH* mRNA. However, the transcriptome Δ*sdsR* bacteria revealed that the abundance of *yeaGH* mRNA did not significantly differ from that in wild-type bacteria (differentially expressed genes were defined as those with a false discovery rate adjusted P < 0.05), suggesting that the increased abundance of YeaG and YeaH in N-24 Δ*sdsR* bacteria is due a post-transcriptional influence of SdsR on *yeaGH* mRNA (Figure S6B).

**Figure 4.**
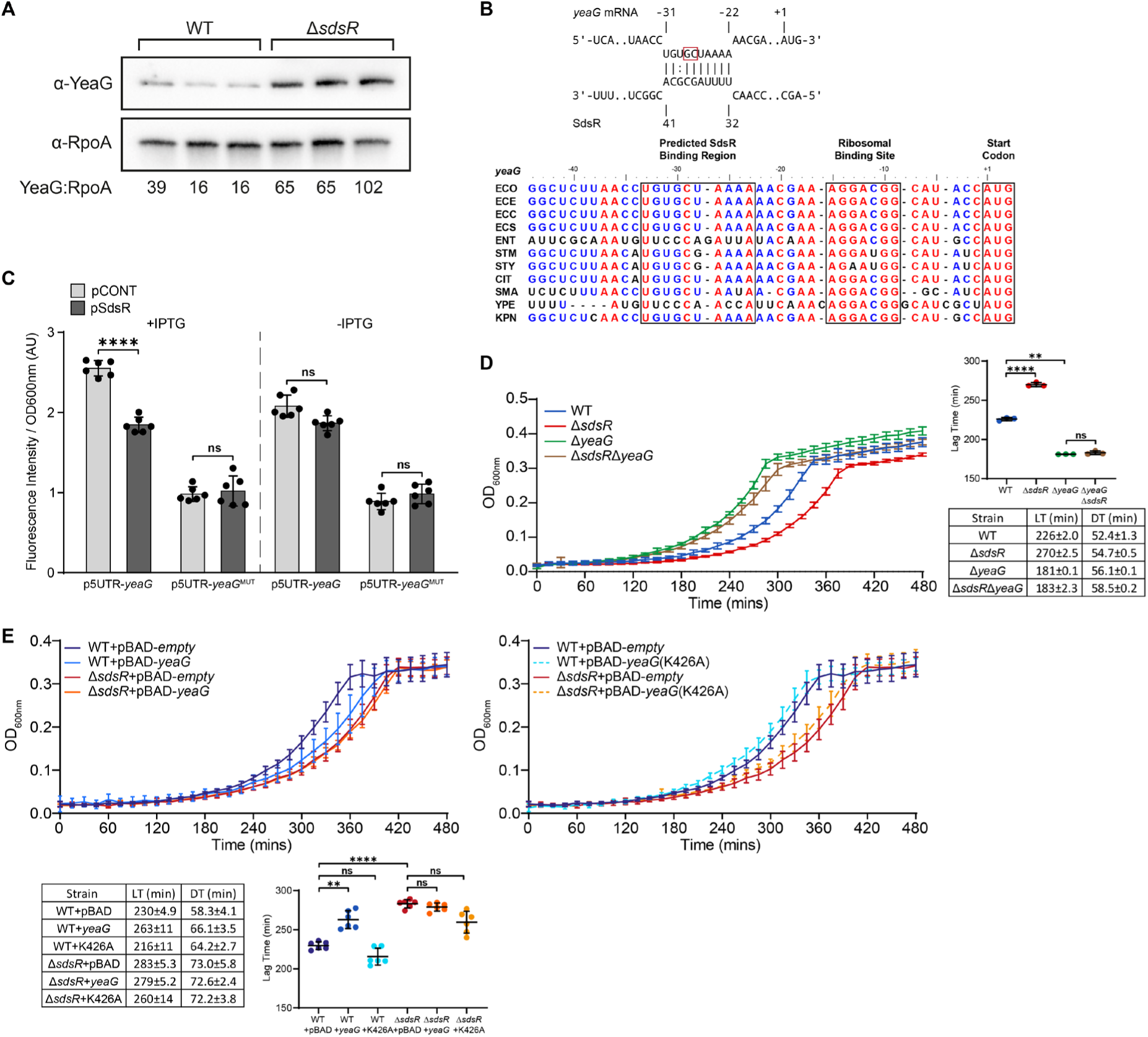
Optimal growth recovery from long-term N starvation requires translational downregulation of the conserved *yeaGH* operon by SdsR. (**A**) Immunoblot of whole-cell extracts of wild-type and Δ*sdsR E. coli*, sampled at N-24, three biological replicates are shown. Probed with anti-YeaG antibody and anti-RpoA antibody (loading control). The ratio of YeaG:RpoA levels are shown below the immunoblot, as determined by Image Lab software. **(B)** Predicted base-pairing interactions between SdsR and the 5’UTR of *yeaG*. The base pairs that were mutated to create *yeaG*^MUT^ are boxed. Predictions were performed using IntraRNA (Bioinformatics Group Freiburg). Alignment of the 5’UTR region of *yeaG* amongst clinically relevant enterobacteria species. Fully, partially, and poorly conserved nucleotides are indicated in red, blue, and black, respectively. The predicted SdsR binding region, RBS and AUG are boxed. Abbreviations correspond to the following species: ECO, *Escherichia coli* K12-MG1655; ECE, *Escherichia coli* O157:H7 str. EDL933; ECC, *Escherichia coli* CFT073; ECS, *Escherichia coli* ST131; ENT, *Enterobacter cloacae* ATCC 13047; SFL, *Shigella flexneri* str. 301; STM, *Salmonella* Typhimurium LT2; STY, *Salmonella typhi* CT18; CIT, *Citrobacter freundii* CFNIH1; SMA, *Serratia marcescens* Db11; YPE, *Yersinia pestis* D182038; KPN, *Klebsiella pneumoniae* HS11286. **(C)** GFP-translational reporter activity in Δ*sdsR* bacteria grown for 10 h in lysogeny broth. Bacteria carried plasmids constitutively expressing the wild-type (p5UTR-*yeaG*) or mutant (p5UTR-*yeaG^MUT^*) 5’UTR of *yeaG* amino-terminally fused to GFP and inducible plasmids either containing SdsR (pSdsR) or a ∼50nt nonsense RNA fragment (pCONT). Values shown are fluorescence intensity normalised to OD_600nm_ both in the presence and absence of IPTG to induced SdsR expression. Error bars represent standard deviation (n=6). Statistical analysis performed by Brown-Forsyth and Welch’s ANOVA. **(D)** Recovery growth of wild-type, Δ*sdsR*, Δ*yeaG* and Δ*sdsR*Δ*yeaG* N-24 bacteria following sub-culturing into fresh culture media. Error bars represent standard deviation (n=3). Additional graph shows lag-time and table shows lag-time (LT) and doubling-time (DT). Statistical analysis performed by Brown-Forsyth and Welch’s ANOVA. **(E)** Recovery growth of N-24 wild-type and Δ*sdsR* bacteria overexpressing either wild-type (pBAD18-*yeaG*) or a catalytically defective mutant (pBAD18-*yeaG* K426A) YeaG following sub-culturing of 24 h N-starved bacteria into fresh culture media. Data is split onto two axis for clarity, but was performed at the same time. Expression of *yeaG* was induced with 0.125% (v/v) L-arabinose at N-. Error bars represent standard deviation (n=6). Additional graph shows lag-time and table shows lag-time (LT) and doubling-time (DT). Statistical analysis performed by Brown-Forsyth and Welch’s ANOVA. (****, P<0.0001; **, P<0.01).

Therefore, to independently validate the SdsR-mediated translational downregulation of *yeaGH*, we used a GFP reporter system in which SdsR and its target sequence were co-expressed from different compatible plasmids (28). As the predicted base-pairing interactions between SdsR and 5’ UTR of *yeaG* mRNA occur between positions -31 and -22 relative to the translation start site (AUG; A is +1; Figure 4B), we fused the nucleotide sequence comprising positions -93 to +63 to the amino terminus of GFP and placed the fusion construct under a constitutive promoter in pXG10 (28) to create p5UTR-*yeaG*. The SdsR sequence was placed under an IPTG-inducible promoter in pKF68 (29) to create plasmid pSdsR. Plasmid pJV300, which expresses a ∼50 nucleotide nonsense RNA derived from *rrnB* terminator region (28), served as the control vector for pSdsR (hereafter referred to as pCONT). As shown in Figure 4C, induction of SdsR expression with IPTG in stationary phase *E. coli* grown in lysogeny broth resulted in a decrease in GFP signal by ∼27% compared to bacteria containing pCONT. This decrease was not seen in control cultures when IPTG was not present, although, we note that overall GFP signal is lower in cultures without IPTG than those with (Figure 4C). We created p5UTR-*yeaG*^MUT^ in which the SdsR interaction sequence in the 5’ UTR was altered to compromise efficient SdsR binding (Figure 4B). As expected, we did not detect the decrease in GFP signal upon expression of SdsR (Figure 4C). We note that the overall GFP signal is lower from p5UTR-*yea*G^MUT^ and suggest that this could be due to the mutations adversely affecting adjacent RBS sequence. In sum, the reporter system independently confirms that RIL-seq and proteomics data and supports our prediction that *yeaGH* mRNA translation is regulated by SdsR.

Next, we considered whether the increased lag time to growth recovery from N-24 displayed by the Δ*sdsR* mutant (Figure 3A) would be absent if YeaG could not be overexpressed. Hence, we created a Δ*sdsR*Δ*yeaG* double mutant *E. coli* strain and compared its lag time to growth recovery form N-24 to that of the Δ*yeaG* and Δ*sdsR* single mutant and wild-type bacteria. As previously reported by us (44), the Δ*yeaG* bacteria displayed a decreased lag time (by ∼47 min) to growth recovery from N-24 compared to wild-type bacteria (Figure 4D, compare green and blue lines, respectively). However, the increased lag time to growth recovery from N-24 displayed by the Δ*sdsR* mutant (relative to wild-type bacteria) was barely present in the Δ*sdsR*Δ*yeaG* double mutant relative to Δ*yeaG* bacteria (Figure 4D, compare brown and green lines, respectively).

Finally, we considered that overexpression of YeaG from an L-arabinose inducible plasmid (pBAD-*yeaG*), in which the native 5’ UTR of *yeaG* is absent and SdsR thus cannot translationally downregulate its expression, would mimic a scenario analogous to the one in Δ*sdsR* bacteria resulting in an increased lag time to growth-recovery from N-24. We grew wild-type bacteria containing pBAD-*yeaG* and pBAD-*empty* to N- and induced YeaG expression with L-arabinose. As expected, the lag time to growth recovery from N-24 wild-type bacteria containing pBAD-*yeaG* was increased by ∼28 min compared to that of bacteria containing pBAD-empty (Figure 4E left, blue and dark blue lines, respectively) and resembled that of Δ*sdsR* bacteria containing pBAD-empty (Figure 4E left, red line). The lag time to growth recovery, when YeaG was overexpressed in Δ*sdsR* bacteria, mirrored that of Δ*sdsR* bacteria containing pBAD-empty (Figure 4E, right, orange line). It seems that overexpressing YeaG does not further increase the lag time to growth recovery of the Δ*sdsR* bacteria. To exclude the possibility that the increased lag-time to growth-recovery of wild-type bacteria containing pBAD-*yeaG* was not due to YeaG overexpression *per se* but instead due to the biological activity of YeaG, we overexpressed a catalytically deleterious YeaG variant containing a K426A mutation in the kinase domain of YeaG (44). As expected, the lag-time to growth-recovery was absent in wild-type and Δ*sdsR* bacteria containing pBAD-*yeaG*(K426A) (Figure 4E right, compare dark blue and red lines with orange dashed and cyan dashed lines, respectively). In sum, we have uncovered that the translational downregulation of the conserved operon *yeaGH* by SdsR is required for optimal growth recovery of long-term N starved *E. coli*.

## DISCUSSION

The past few years have seen a surge in new methodologies to identify post-transcriptional regulatory networks in bacteria at a global scale, which have underscored the depth and breadth of their contribution to gene expression (58). However, our current knowledge of post-transcriptional regulatory networks is restricted to growing bacteria, bacteria undergoing growth transitions (e.g., from the exponential to stationary phase of growth) or have been exposed to iron limitation (4,18,21). The impetus for this study was to understand to what extent the Hfq-mediated post-transcriptional regulatory network, i.e., the RNA-RNA interactome, changes during growth arrest and underpins recovery from growth arrest. In addition, as we used N starvation to induce growth arrest, this allowed us to simultaneously gain deeper insights into the post-transcriptional basis of the adaptive response to N starvation in *E. coli*.

Our results have revealed that the Hfq-mediated RNA-RNA interactome undergoes large scale reprogramming, not only between growth and growth arrested states (i.e., N+ to N- and N-24 to N-24+2 states), but also throughout the course of growth arrest (i.e., N- to N-24 state) (Figure 1). One way this reprogramming manifests itself is via the different sRNA that govern the Hfq-mediated RNA-RNA interactome at different growth states. For instance, we detected SdsR in ∼30% of chimeras in the Hfq-mediated RNA-RNA interactome of N-24 bacteria. However, it is only present in less than 5% of chimeras in the Hfq-mediated RNA-RNA interactome of bacteria from N-, and in less than 1% of chimeras in bacteria from N+ and N-24+2 (Figure 2A & Figure S7). This is consistent with previous RIL-seq results showing increased prominence of SdsR in the Hfq-mediated RNA-RNA interactome of bacteria from the stationary phase of growth (18). Conversely, ArcZ is an sRNA that governs the Hfq-mediated RNA-RNA interactome at all growth states studied here (Figure 2A), but its target suite differs substantially between growth states (Figure S7), with only 11% of its targets being found in its chimeras under all growth states. Notably, the largest proportion (∼37%) of all chimeras involving ArcZ in Hfq-mediated RNA-RNA interactome in N-bacteria is with the sRNA RybB, suggesting that regulation of sRNA activity by sponging interactions can be growth state specific phenomenon. In contrast, we identified a number of RNA-RNA interactions that were detected across all growth states - we collectively referred to this as the ‘core’ Hfq-mediated RNA-RNA interactome (Figure 1G). Although, the number of chimeras that constitute the core Hfq-mediated RNA-RNA interactome unsurprisingly fluctuate between the four growth conditions used here (Figure S1E), which underscores the inherent plasticity of post-transcriptional regulatory networks in bacteria. Notably, 49% of the core Hfq-mediated RNA-RNA interactions identified under our conditions were also detected in *E. coli* during exponential growth and in the early stationary phase in lysogeny broth (18) (Figure S8) suggesting that a portion of the Hfq-mediated post-transcriptional regulatory network may act in a ‘housekeeping’ capacity, rather than in a condition specific regulatory capacity.

Conversely, the RNA-RNA interactions identified in growing bacteria in this study (i.e., at N+ and N-24+2 states) also differs from the Hfq-mediated RNA-RNA interactome identified by Melamed and colleagues in exponentially growing *E. coli* in lysogeny broth (18). For example, of the 315 and 279 unique RNA-RNA interactions identified in the Hfq-mediated RNA interactome of N+ and N-24+2 bacteria, respectively, only 110 and 84 of them are also detected in the Hfq-mediated RNA-RNA interactome of exponentially growing *E. coli* in lysogeny broth. This implies that the Hfq-mediated RNA-RNA interactome is highly sensitive to changes in growth conditions even if they permit active growth. Consistent with this view, the Hfq-mediated RNA-RNA interactomes of N+ and N-24+2 bacteria are similar but not identical (Figure 1C and Figure 1D). Overall, the sensitivity and specificity to growth conditions may underpin the inherent physiological purpose of post-transcriptional regulatory networks, where they might be considered to fine tune (see below), rather than to act in a binary fashion, to regulate the flow of genetic information to meet cellular needs.

Our study also suggests how a single sRNA can couple two regulons to potentially fine tune an adaptive response. This is exemplified by GlnZ whose expression is activated by NtrC and is thus a member of the NtrC regulon. The regulatory targets of GlnZ have been explored in a number of previous studies (50,51), and many of those identified previously have also been identified in the currently study (Figure S3A). However, our data additionally uncovered that GlnZ interacts with the mRNA of *nac* in the N-growth state, which encodes a global transcription factor and is a member of the NtrC regulon. The interaction site of GlnZ is in the CDS of *nac* mRNA (Figure S9) and at this stage we are uncertain whether GlnZ has a positive or negative regulatory effect on Nac expression. The interaction between GlnZ and *nac* mRNA is only detected in the N-state and it is absent in the N-24 state (Figure S3A). Thus, we propose that GlnZ functions to adjust Nac proteins levels to fine tune the adaptive response to N starvation as N starvation condition ensues. For example, GlnZ could have a positive regulatory effect on *nac* mRNA and thereby increase Nac proteins levels in N-bacteria to repress the transcription of *sucA*. As *sucA* encodes a subunit of 2-oxoglutarate dehydrogenase enzyme, the cooperative action of GlnZ and Nac could allow efficient balancing of carbon and nitrogen metabolism at the onset of N starvation (50,51). Further, our analysis of RNA-RNA chimeras involving NtrC activated genes from different stages of N starvation in *E. coli* underscores that post-transcriptional regulation is clearly an important, yet relatively underexplored, facet of the adaptive response to N starvation in *E. coli* (Figure S3B).

Although many of the new global approaches provide a comprehensive overview of post-transcriptional regulatory interactions at the transcriptome wide level, the major challenge for many investigators is to ascertain their impact on gene expression and ultimately cellular physiology. This is further compounded by the fact that deletion of an sRNA of interest often does not result in discernible phenotypic changes. We have demonstrated that coupling RIL-seq with global comparative proteomics of wild-type and a deletion mutant of an sRNA of interest can not only validate whether the interactions made by the said sRNA are productive, i.e., lead to changes in gene expression, but also differentiate between the direct and indirect effects of how the sRNA contributes to gene expression. We have shown that the absence of SdsR, the sRNA that governs the Hfq-mediated RNA-RNA interactome in N-24 bacteria, results in a large scale perturbation of the proteome of *E. coli*, extending far beyond its direct regulatory targets (Figure 2C). However, this interpretation must be viewed cautiously as the observed large scale changes to the proteome could be the accumulative consequence of dysregulated post-transcriptional events that occurred in the N+ and N-growth states. Nonetheless, at N-24, we observed that the expected regulatory outcomes of SdsR targeted mRNAs correlated surprisingly well with the levels of proteins encodes by the targeted mRNAs (Figure 2D). The target spectrum of SdsR in N-24 bacteria is functionally diverse, containing genes associated with almost every major cellular function (Figure S5). We noted an enrichment of targets involved in energy metabolism (e.g., *gltA* and *aceA*) carbon metabolism (e.g., *tktA* and *suhB*) and nitrogen metabolism (e.g., *aroF*, *thrA, serC, solA, asd*) amongst those targets as potentially positively regulated by SdsR (Figure S5 and Figure S10). This implies a role for SdsR in regulating cellular metabolism in N-24 bacteria. This view is consistent with SdsR regulating a number of targets associated with energy metabolism in stationary phase *Salmonella* bacteria (29).

We also observed a number of proteins (TolC, TamA, MepK, FumC) associated with antibiotic susceptibility are upregulated in the proteome of Δ*sdsR* bacteria (Figure S10), suggesting that SdsR acts to post-transcriptionally downregulate the expression of them. Consistent with this view, Parker and Gottesman (59) previously reported that SdsR downregulates expression of the outer membrane channel TolC, and that this has a subtle effect on susceptibility to novobiocin and erythromycin. Further, mRNAs of genes encoding TamA, a component of the TAM translocation and assembly module that has been linked to antimicrobial stress in *K. pneumoniae* (60), MepK, a regulator of alternate peptidoglycan biosynthesis that has been linked to beta-lactam resistance (61,62) and FumC, the alternative fumarase that has been linked to susceptibility of *E. coli* to certain bactericidal antibiotics (63) also belong to the regulon of SdsR and their expression is potentially negatively affected by SdsR. In contrast, the D-ala-D-ala ligase DdlB, which is directly targeted by the antibiotic cycloserine (64), was downregulated in the proteome of Δ*sdsR* bacteria (Figure S10), suggesting that SdsR acts to post-transcriptionally upregulate DdlB expression. Overall, our results support the widening body of evidence that sRNAs, and the RNA-RNA interactomes defined by them, may contribute to modulating the antibiotic susceptibility of bacteria (reviewed in (65)).

Despite the broad involvement of SdsR in diverse cellular processes in growth arrested bacteria (Figure S10 & (29)), under N starvation, the absence of SdsR, surprisingly, neither effects growth nor viability. However, we discovered a role for SdsR in affecting growth recovery from long-term N starvation and uncovered that this process is underpinned by two new members, *yeaG* and *yeaH*, of the SdsR regulon in N-24 bacteria. In previous work, we identified the highly conserved *yeaGH* operon as part of the NtrC regulon and showed that YeaG and YeaH function together to quiesce metabolism upon sensing N starvation (66). Therefore, the absence of YeaG and YeaH are detrimental to cell viability as a function of time under N starvation, but conversely Δ*yeaG,* Δ*yeaH* and Δ*yeaGH* bacteria recover growth faster from N starvation following inoculation into fresh media. Put simply, YeaG and YeaH can be considered to function as a metabolic brake. YeaG protein levels peak during the first 9 h into N starvation but subsequently drops as N starvation ensues – perhaps suggesting that YeaG is only required in the early stages of N starvation to quiesce metabolism. Although the mechanism by which YeaG and YeaH induce metabolic quiescence will be a topic of a future study, the discovery that *yeaG* mRNA is subjected to post-transcriptional regulation by SdsR in long-term N starved bacteria (Figure 2D & Figure 4C) underscores the importance of controlling YeaG (and YeaH) levels as N starvation ensues. The management of YeaG (and YeaH) levels seems important to balance quiescing metabolism under growth non-permissive conditions and to restart metabolism to allow optimal growth recovery when conditions become growth permissive. Consistent with this view, failure to do so adversely effects the ability of bacteria to optimally recover from long-term N starvation induced growth arrest.

In sum, our study has underscored the importance of studying RNA interactomes of growth arrested bacteria as they can be dynamic and extensive as those in actively growing bacteria and, importantly, can contribute to uncovering new biological processes that govern bacterial growth. As the *yeaGH* operon and SdsR are conserved in many bacteria, including ESKAPE pathogens (Figure 4A & Figure S4), our study has uncovered a conserved post-transcriptional regulatory axis that governs optimal growth recovery from long-term N starvation. We speculate that the *yeaGH*-SdsR regulatory axis might contribute to optimal growth recovery from diverse conditions that induce growth arrest and thus identifying ways to interfere with this regulatory axis might provide options to manage bacterial growth and colonisation. Similarly, the core Hfq-mediated RNA-RNA interactome offers several opportunities for antibacterial intervention. In this regard, our study provides a treasure chest of RNA-RNA interactions in a prototypical bacterium at four different states of growth that can be exploited to gain valuable new insights into the post-transcriptional regulatory basis of bacterial growth and growth arrest.

## DATA AVAILABILITY

The RNA-seq and RIL-seq data discussed in this publication are accessible through ArrayExpress E-MTAB-13402 and E-MTAB-13403 respectively. The proteomics data can be accessed through PRIDE PXD045656.

## REFERENCES

1. Vogel, J. and Luisi, B.F. (2011) Hfq and its constellation of RNA. Nat Rev Microbiol, 9, 578–589.

2. Kavita, K., de Mets, F. and Gottesman, S. (2018) New aspects of RNA-based regulation by Hfq and its partner sRNAs. Current opinion in microbiology, 42, 53–61.

3. Olejniczak, M. and Storz, G. (2017) ProQ/FinO-domain proteins: another ubiquitous family of RNA matchmakers? Molecular microbiology, 104, 905–915.

4. Melamed, S., Adams, P.P., Zhang, A., Zhang, H. and Storz, G. (2020) RNA-RNA Interactomes of ProQ and Hfq Reveal Overlapping and Competing Roles. Mol Cell, 77, 411–425 e417.

5. Papenfort, K. and Melamed, S. (2023) Small RNAs, Large Networks: Posttranscriptional Regulons in Gram-Negative Bacteria. Annu Rev Microbiol.

6. Kavita, K., de Mets, F. and Gottesman, S. (2018) New aspects of RNA-based regulation by Hfq and its partner sRNAs. Current opinion in microbiology, 42, 53–61.

7. Frohlich, K.S. and Papenfort, K. (2020) Regulation outside the box: New mechanisms for small RNAs. Mol Microbiol, 114, 363–366.

8. Majdalani, N., Cunning, C., Sledjeski, D., Elliott, T. and Gottesman, S. (1998) DsrA RNA regulates translation of RpoS message by an anti-antisense mechanism, independent of its action as an antisilencer of transcription. Proc Natl Acad Sci U S A, 95, 12462–12467.

9. Beisel, C.L., Updegrove, T.B., Janson, B.J. and Storz, G. (2012) Multiple factors dictate target selection by Hfq-binding small RNAs. EMBO J, 31, 1961–1974.

10. Frohlich, K.S., Papenfort, K., Fekete, A. and Vogel, J. (2013) A small RNA activates CFA synthase by isoform-specific mRNA stabilization. EMBO J, 32, 2963–2979.

11. Papenfort, K., Sun, Y., Miyakoshi, M., Vanderpool, C.K. and Vogel, J. (2013) Small RNA-mediated activation of sugar phosphatase mRNA regulates glucose homeostasis. Cell, 153, 426–437.

12. Prevost, K., Desnoyers, G., Jacques, J.F., Lavoie, F. and Masse, E. (2011) Small RNA-induced mRNA degradation achieved through both translation block and activated cleavage. Genes Dev, 25, 385–396.

13. Denham, E.L. (2020) The Sponge RNAs of bacteria - How to find them and their role in regulating the post-transcriptional network. Biochim Biophys Acta Gene Regul Mech, 1863, 194565.

14. Fuchs, M., Lamm-Schmidt, V., Lence, T., Sulzer, J., Bublitz, A., Wackenreuter, J., Gerovac, M., Strowig, T. and Faber, F. (2023) A network of small RNAs regulates sporulation initiation in Clostridioides difficile. EMBO J, e112858.

15. Miyakoshi, M., Okayama, H., Lejars, M., Kanda, T., Tanaka, Y., Itaya, K., Okuno, M., Itoh, T., Iwai, N. and Wachi, M. (2022) Mining RNA-seq data reveals the massive regulon of GcvB small RNA and its physiological significance in maintaining amino acid homeostasis in Escherichia coli. Mol Microbiol, 117, 160–178.

16. Matera, G., Altuvia, Y., Gerovac, M., El Mouali, Y., Margalit, H. and Vogel, J. (2022) Global RNA interactome of Salmonella discovers a 5’ UTR sponge for the MicF small RNA that connects membrane permeability to transport capacity. Mol Cell, 82, 629–644 e624.

17. Huber, M., Lippegaus, A., Melamed, S., Siemers, M., Wucher, B.R., Hoyos, M., Nadell, C., Storz, G. and Papenfort, K. (2022) An RNA sponge controls quorum sensing dynamics and biofilm formation in Vibrio cholerae. Nat Commun, 13, 7585.

18. Melamed, S., Peer, A., Faigenbaum-Romm, R., Gatt, Y.E., Reiss, N., Bar, A., Altuvia, Y., Argaman, L. and Margalit, H. (2016) Global Mapping of Small RNA-Target Interactions in Bacteria. Mol Cell, 63, 884–897.

19. Faigenbaum-Romm, R., Reich, A., Gatt, Y.E., Barsheshet, M., Argaman, L. and Margalit, H. (2020) Hierarchy in Hfq Chaperon Occupancy of Small RNA Targets Plays a Major Role in Their Regulation. Cell Rep, 30, 3127–3138 e3126.

20. Pearl Mizrahi, S., Elbaz, N., Argaman, L., Altuvia, Y., Katsowich, N., Socol, Y., Bar, A., Rosenshine, I. and Margalit, H. (2021) The impact of Hfq-mediated sRNA-mRNA interactome on the virulence of enteropathogenic Escherichia coli. Sci Adv, 7, eabi8228.

21. Iosub, I.A., van Nues, R.W., McKellar, S.W., Nieken, K.J., Marchioreto, M., Sy, B., Tree, J.J., Viero, G. and Granneman, S. (2020) Hfq CLASH uncovers sRNA-target interaction networks linked to nutrient availability adaptation. Elife, 9.

22. Gebhardt, M.J., Farland, E.A., Basu, P., Macareno, K., Melamed, S. and Dove, S.L. (2023) Hfq-licensed RNA-RNA interactome in Pseudomonas aeruginosa reveals a keystone sRNA. Proc Natl Acad Sci U S A, 120, e2218407120.

23. Melamed, S., Faigenbaum-Romm, R., Peer, A., Reiss, N., Shechter, O., Bar, A., Altuvia, Y., Argaman, L. and Margalit, H. (2018) Mapping the small RNA interactome in bacteria using RIL-seq. Nat Protoc, 13, 1–33.

24. Reese, A.T., Pereira, F.C., Schintlmeister, A., Berry, D., Wagner, M., Hale, L.P., Wu, A., Jiang, S., Durand, H.K., Zhou, X. et al. (2018) Microbial nitrogen limitation in the mammalian large intestine. Nat Microbiol, 3, 1441–1450.

25. Elser, J.J., Bracken, M.E., Cleland, E.E., Gruner, D.S., Harpole, W.S., Hillebrand, H., Ngai, J.T., Seabloom, E.W., Shurin, J.B. and Smith, J.E. (2007) Global analysis of nitrogen and phosphorus limitation of primary producers in freshwater, marine and terrestrial ecosystems. Ecol Lett, 10, 1135–1142.

26. Zimmer, D.P., Soupene, E., Lee, H.L., Wendisch, V.F., Khodursky, A.B., Peter, B.J., Bender, R.A. and Kustu, S. (2000) Nitrogen regulatory protein C-controlled genes of Escherichia coli: scavenging as a defense against nitrogen limitation. Proc Natl Acad Sci U S A, 97, 14674–14679.

27. Brown, D.R., Barton, G., Pan, Z., Buck, M. and Wigneshweraraj, S. (2014) Nitrogen stress response and stringent response are coupled in Escherichia coli. Nat Commun, 5, 4115.

28. Urban, J.H. and Vogel, J. (2007) Translational control and target recognition by Escherichia coli small RNAs in vivo. Nucleic Acids Res, 35, 1018–1037.

29. Frohlich, K.S., Haneke, K., Papenfort, K. and Vogel, J. (2016) The target spectrum of SdsR small RNA in Salmonella. Nucleic Acids Res, 44, 10406–10422.

30. Gibson, D.G., Young, L., Chuang, R.Y., Venter, J.C., Hutchison, C.A., 3rd and Smith, H.O. (2009) Enzymatic assembly of DNA molecules up to several hundred kilobases. Nature methods, 6, 343-345.

31. Atlas, R.M. (2010) Handbook of Microbiological Media, Fourth Edition. CRC Press.

32. Mann, M., Wright, P.R. and Backofen, R. (2017) IntaRNA 2.0: enhanced and customizable prediction of RNA-RNA interactions. Nucleic Acids Res, 45, W435–W439.

33. Hughes, C.S., Foehr, S., Garfield, D.A., Furlong, E.E., Steinmetz, L.M. and Krijgsveld, J. (2014) Ultrasensitive proteome analysis using paramagnetic bead technology. Mol Syst Biol, 10, 757.

34. Moggridge, S., Sorensen, P.H., Morin, G.B. and Hughes, C.S. (2018) Extending the Compatibility of the SP3 Paramagnetic Bead Processing Approach for Proteomics. J Proteome Res, 17, 1730–1740.

35. Kong, A.T., Leprevost, F.V., Avtonomov, D.M., Mellacheruvu, D. and Nesvizhskii, A.I. (2017) MSFragger: ultrafast and comprehensive peptide identification in mass spectrometry-based proteomics. Nature methods, 14, 513–520.

36. Savitski, M.M., Wilhelm, M., Hahne, H., Kuster, B. and Bantscheff, M. (2015) A Scalable Approach for Protein False Discovery Rate Estimation in Large Proteomic Data Sets. Molecular & cellular proteomics: MCP, 14, 2394–2404.

37. Ritchie, M.E., Phipson, B., Wu, D., Hu, Y., Law, C.W., Shi, W. and Smyth, G.K. (2015) limma powers differential expression analyses for RNA-sequencing and microarray studies. Nucleic Acids Res, 43, e47.

38. Huber, W., von Heydebreck, A., Sultmann, H., Poustka, A. and Vingron, M. (2002) Variance stabilization applied to microarray data calibration and to the quantification of differential expression. Bioinformatics, 18 Suppl 1, S96–104.

39. Switzer, A., Evangelopoulos, D., Figueira, R., de Carvalho, L.P.S., Brown, D.R. and Wigneshweraraj, S. (2018) A novel regulatory factor affecting the transcription of methionine biosynthesis genes in Escherichia coli experiencing sustained nitrogen starvation. Microbiology (Reading, England).

40. Levin-Reisman, I., Fridman, O. and Balaban, N.Q. (2014) ScanLag: high-throughput quantification of colony growth and lag time. J Vis Exp.

41. Stead, M.B., Agrawal, A., Bowden, K.E., Nasir, R., Mohanty, B.K., Meagher, R.B. and Kushner, S.R. (2012) RNAsnap: a rapid, quantitative and inexpensive, method for isolating total RNA from bacteria. Nucleic Acids Res, 40, e156.

42. Love, M.I., Huber, W. and Anders, S. (2014) Moderated estimation of fold change and dispersion for RNA-seq data with DESeq2. Genome Biol, 15, 550.

43. Switzer, A., Burchell, L., McQuail, J. and Wigneshweraraj, S. (2020) The Adaptive Response to Long-Term Nitrogen Starvation in Escherichia coli Requires the Breakdown of Allantoin. J Bacteriol, 202.

44. Figueira, R., Brown, D.R., Ferreira, D., Eldridge, M.J., Burchell, L., Pan, Z., Helaine, S. and Wigneshweraraj, S. (2015) Adaptation to sustained nitrogen starvation by Escherichia coli requires the eukaryote-like serine/threonine kinase YeaG. Sci Rep, 5, 17524.

45. Switzer, A., Evangelopoulos, D., Figueira, R., de Carvalho, L.P.S., Brown, D.R. and Wigneshweraraj, S. (2018) A novel regulatory factor affecting the transcription of methionine biosynthesis genes in Escherichia coli experiencing sustained nitrogen starvation. Microbiology (Reading), 164, 1457–1470.

46. Hor, J., Matera, G., Vogel, J., Gottesman, S. and Storz, G. (2020) Trans-Acting Small RNAs and Their Effects on Gene Expression in Escherichia coli and Salmonella enterica. EcoSal Plus, 9.

47. Burtnick, M.N., Brett, P.J., Harding, S.V., Ngugi, S.A., Ribot, W.J., Chantratita, N., Scorpio, A., Milne, T.S., Dean, R.E., Fritz, D.L., et al. (2011) The cluster 1 type VI secretion system is a major virulence determinant in Burkholderia pseudomallei. Infection and immunity, 79, 1512–1525.

48. Bender, R.A. (2010) A NAC for regulating metabolism: the nitrogen assimilation control protein (NAC) from Klebsiella pneumoniae. J Bacteriol, 192, 4801–4811.

49. Sanchez-Vazquez, P., Dewey, C.N., Kitten, N., Ross, W. and Gourse, R.L. (2019) Genome-wide effects on Escherichia coli transcription from ppGpp binding to its two sites on RNA polymerase. Proc Natl Acad Sci U S A, 116, 8310–8319.

50. Miyakoshi, M., Morita, T., Kobayashi, A., Berger, A., Takahashi, H., Gotoh, Y., Hayashi, T. and Tanaka, K. (2022) Glutamine synthetase mRNA releases sRNA from its 3’UTR to regulate carbon/nitrogen metabolic balance in Enterobacteriaceae. Elife, 11.

51. Walling, L.R., Kouse, A.B., Shabalina, S.A., Zhang, H. and Storz, G. (2022) A 3’ UTR-derived small RNA connecting nitrogen and carbon metabolism in enteric bacteria. Nucleic Acids Res, 50, 10093–10109.

52. Frohlich, K.S., Papenfort, K., Berger, A.A. and Vogel, J. (2012) A conserved RpoS-dependent small RNA controls the synthesis of major porin OmpD. Nucleic Acids Res, 40, 3623–3640.

53. Weber, H., Polen, T., Heuveling, J., Wendisch, V.F. and Hengge, R. (2005) Genome-wide analysis of the general stress response network in Escherichia coli: sigmaS-dependent genes, promoters, and sigma factor selectivity. J Bacteriol, 187, 1591–1603.

54. Rozen, Y. and Belkin, S. (2001) Survival of enteric bacteria in seawater. FEMS Microbiol Rev, 25, 513–529.

55. Gyaneshwar, P., Paliy, O., McAuliffe, J., Popham, D.L., Jordan, M.I. and Kustu, S. (2005) Sulfur and nitrogen limitation in Escherichia coli K-12: specific homeostatic responses. J Bacteriol, 187, 1074–1090.

56. Tagourti, J., Landoulsi, A. and Richarme, G. (2008) Cloning, expression, purification and characterization of the stress kinase YeaG from Escherichia coli. Protein expression and purification, 59, 79–85.

57. Stancik, I.A., Sestak, M.S., Ji, B., Axelson-Fisk, M., Franjevic, D., Jers, C., Domazet-Loso, T. and Mijakovic, I. (2018) Serine/Threonine Protein Kinases from Bacteria, Archaea and Eukarya Share a Common Evolutionary Origin Deeply Rooted in the Tree of Life. J Mol Biol, 430, 27–32.

58. Melamed, S. (2020) New sequencing methodologies reveal interplay between multiple RNA-binding proteins and their RNAs. Curr Genet, 66, 713–717.

59. Parker, A. and Gottesman, S. (2016) Small RNA Regulation of TolC, the Outer Membrane Component of Bacterial Multidrug Transporters. Journal of bacteriology, 198, 1101–1113.

60. Jung, H.J., Sorbara, M.T. and Pamer, E.G. (2021) TAM mediates adaptation of carbapenem-resistant Klebsiella pneumoniae to antimicrobial stress during host colonization and infection. PLoS pathogens, 17, e1009309.

61. Chodiseti, P.K. and Reddy, M. (2019) Peptidoglycan hydrolase of an unusual cross-link cleavage specificity contributes to bacterial cell wall synthesis. Proceedings of the National Academy of Sciences of the United States of America, 116, 7825–7830.

62. Suarez, S.A. and Martiny, A.C. (2021) Gene Amplification Uncovers Large Previously Unrecognized Cryptic Antibiotic Resistance Potential in E. coli. Microbiol Spectr, 9, e0028921.

63. Himpsl, S.D., Shea, A.E., Zora, J., Stocki, J.A., Foreman, D., Alteri, C.J. and Mobley, H.L.T. (2020) The oxidative fumarase FumC is a key contributor for E. coli fitness under iron-limitation and during UTI. PLoS pathogens, 16, e1008382.

64. Neuhaus, F.C. and Lynch, J.L. (1964) The Enzymatic Synthesis of D-Alanyl-D-Alanine. 3. On the Inhibition of D-Alanyl-D-Alanine Synthetase by the Antibiotic D-Cycloserine. Biochemistry, 3, 471–480.

65. Mediati, D.G., Wu, S., Wu, W. and Tree, J.J. (2021) Networks of Resistance: Small RNA Control of Antibiotic Resistance. Trends Genet, 37, 35–45.

66. Figueira, R., Brown, D.R., Ferreira, D., Eldridge, M.J.G., Burchell, L., Pan, Z., Helaine, S. and Wigneshweraraj, S. (2015) Adaptation to sustained nitrogen starvation by Escherichia coli requires the eukaryote-like serine/threonine kinase YeaG. Scientific reports, 5, 17524.

