## Supplementary Figures 1-10, Supplementary Table 1 for "Global RNA interactome of nitrogen starved *Escherichia coli* uncovers a conserved post-transcriptional regulatory axis required for optimal growth recovery"

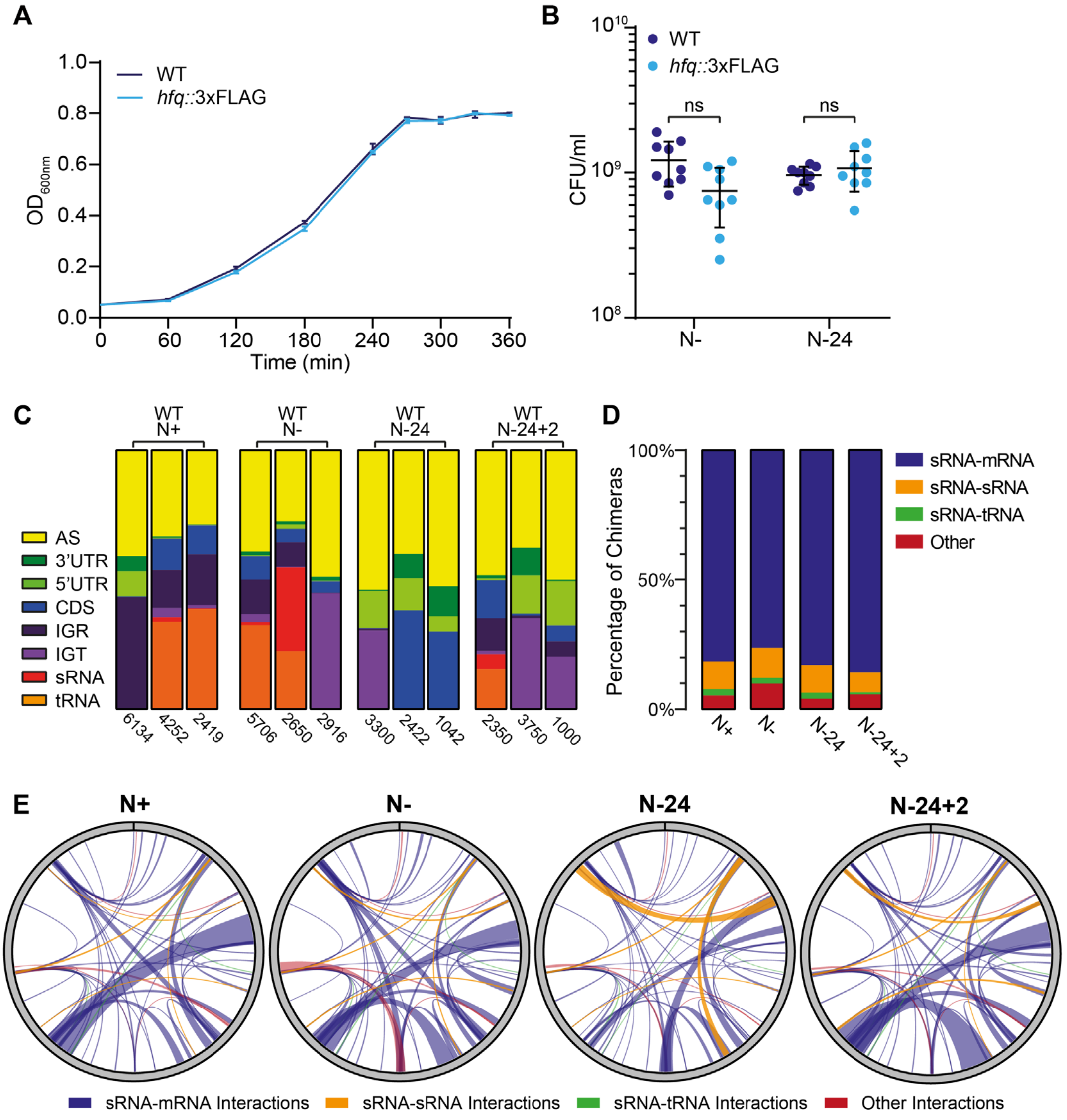


**Figure S1**. **(A)** Growth of wild-type and *hfq-FLAG* bacteria under N limiting conditions. Error bars represent standard deviation (n = 3). **(B)** Viability of wild-type and *hfq-FLAG* bacteria following 20 min (N-) and 24 h (N-24) of N starvation, measured by counting CFUs. Error bars represent standard deviation (n = 9). Statistical analysis performed by Welch's T-tests. **(C)** Relative frequencies of each RNA type found in chimeric fragments, in individual replicates across all time points for the wild-type datasets. **(D)** Relative frequencies of each RNA-RNA interaction type found in chimeric fragments across all time points for *hfq-FLAG* datasets. **(E)** Circos plots of the RIL-seq interactions which are detected in every time point that are represented by at least 30 chimeric fragments in two individual replicates. The thickness of each connection is proportional to the average number of chimeras detected for a given interaction across the three replicates. sRNA:mRNA, sRNA:sRNA, sRNA:tRNA and other interactions are represented by blue, orange, green and red lines respectively.

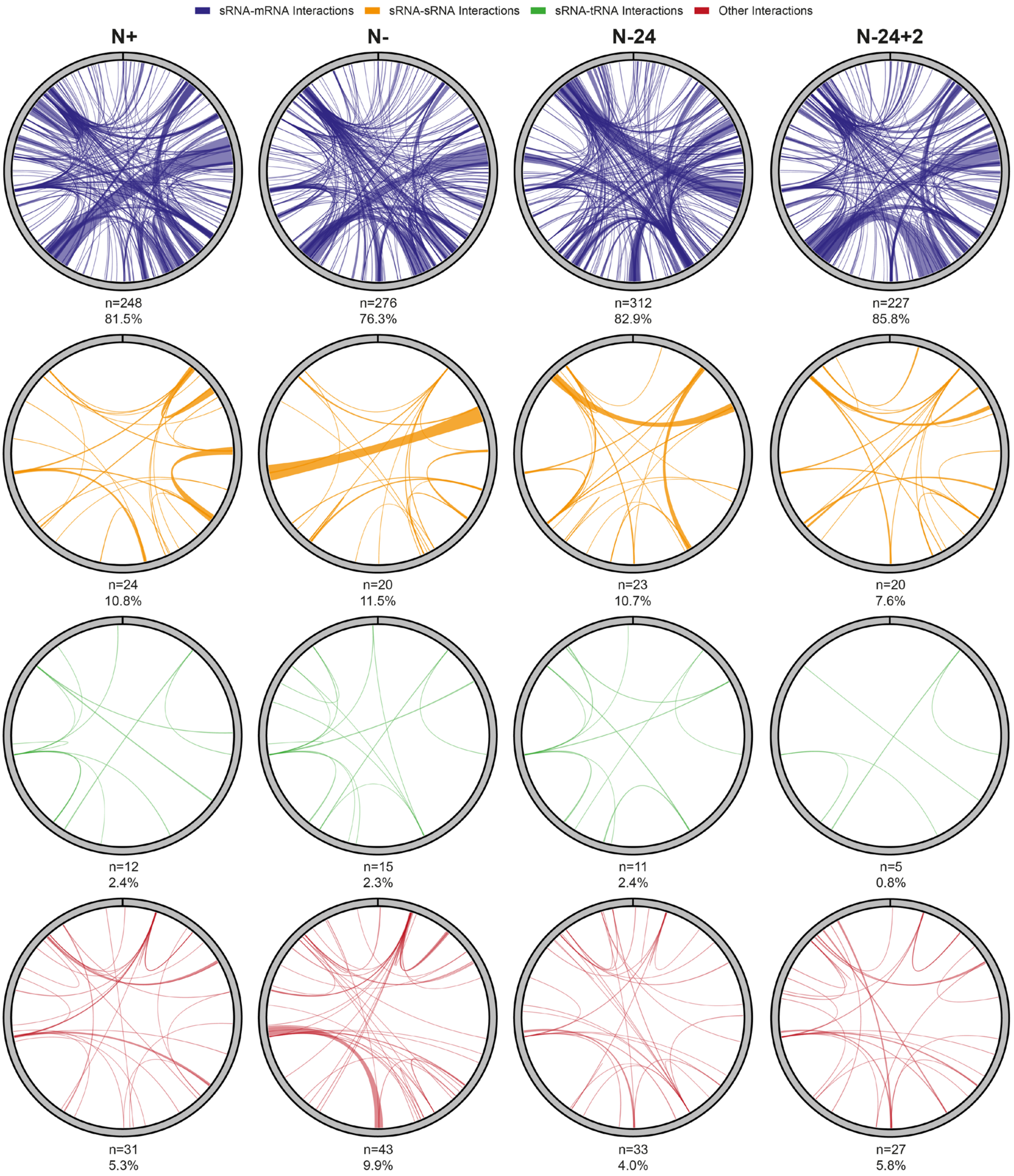


**Figure S2**. Circos plots of sRNA:mRNA, sRNA:sRNA, sRNA:tRNA and other interactions at each time point, that are represented by at least 30 chimeric fragments in two individual replicates. The thickness of each connection is proportional to the average number of chimeras detected for a given interaction across the three replicates. The number of interactions of each type, and the respective proportion of the total interactome at that time-point are shown below each circus plot.

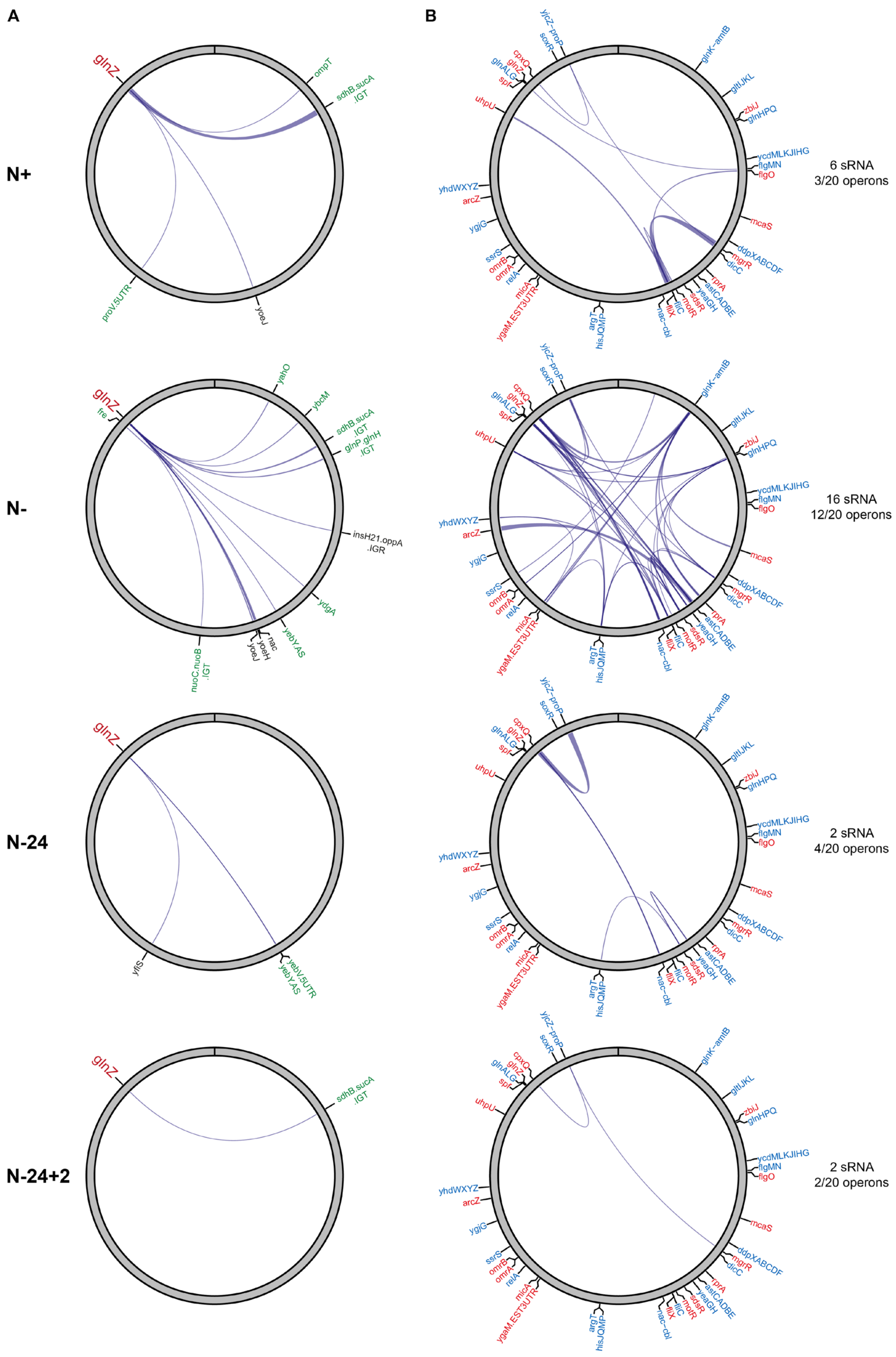


**Figure S3**. **(A)** Circos plots of interactions involving GlnZ that are represented by at least 30 chimeric fragments in two individual replicates at each time point. The thickness of each connection is proportional to the average number of chimeras detected for a given interaction across the three replicates. The targets of GlnZ previously identified in MAPS experiments (1) are shown in green. **(B)** Circos plots of interactions involving mRNA of genes that belong to the Ntr regulon (i.e., genes directed activated by NtrC), at each time point. NtrC dependent operons are shown in blue, and sRNAs are shown in red. The number of potentially regulated operons, and the number of involved sRNA are indicated for each time point.

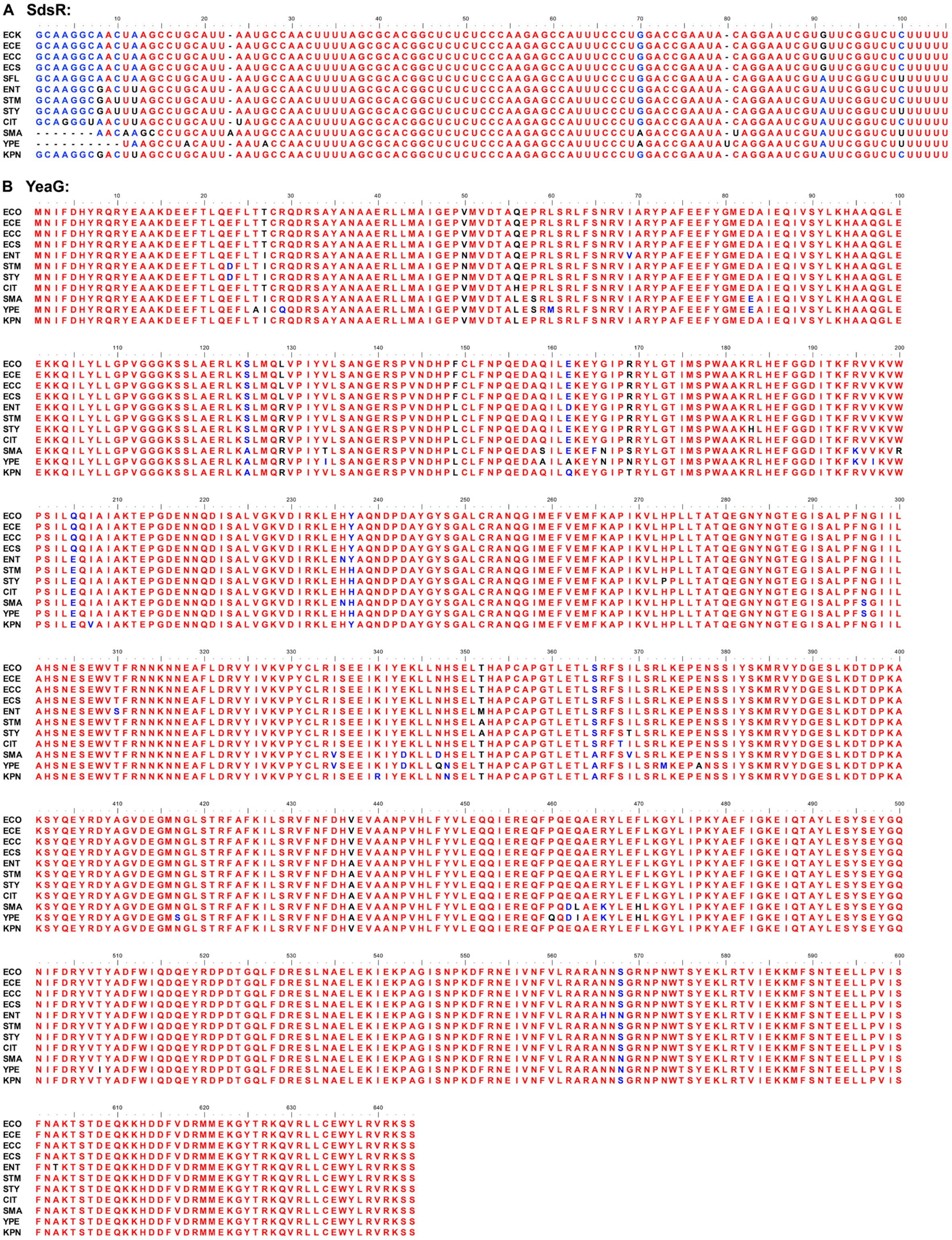
A SdsR:
B YeaG:

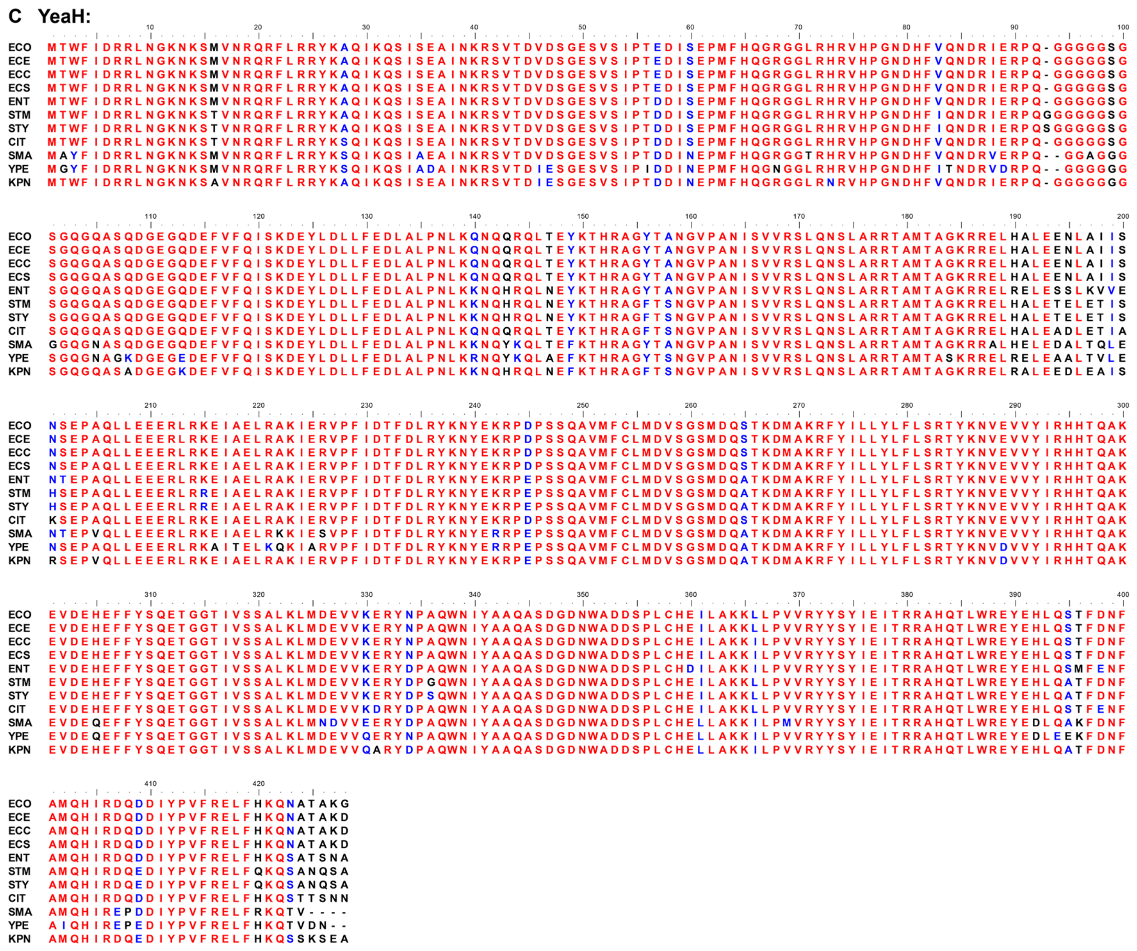

**Figure S4**. **(A)** Alignment of SdsR amongst clinically relevant enterobacteria species. Fully, partially, and poorly conserved nucleotides are indicated in red, blue, and black, respectively. Abbreviations correspond to the following species: ECO, *Escherichia coli* K12-MG1655; ECE, *Escherichia coli* O157:H7 str. EDL933; ECC, *Escherichia coli* CFT073; ECS, *Escherichia coli* ST131; ENT, *Enterobacter cloacae* ATCC 13047; SFL, *Shigella flexneri* str. 301; STM, *Salmonella* Typhimurium LT2; STY, *Salmonella typhi* CT18; CIT, *Citrobacter freundii* CFNIH1; SMA, *Serratia marcescens* Db11; YPE, *Yersinia pestis* D182038; KPN, *Klebsiella pneumoniae* HS11286. **(B, C)** Alignment of the protein sequence of YeaG and YeaH amongst clinically relevant enterobacteria species. Fully, partially, and poorly conserved amino acids are indicated in red, blue, and black, respectively.

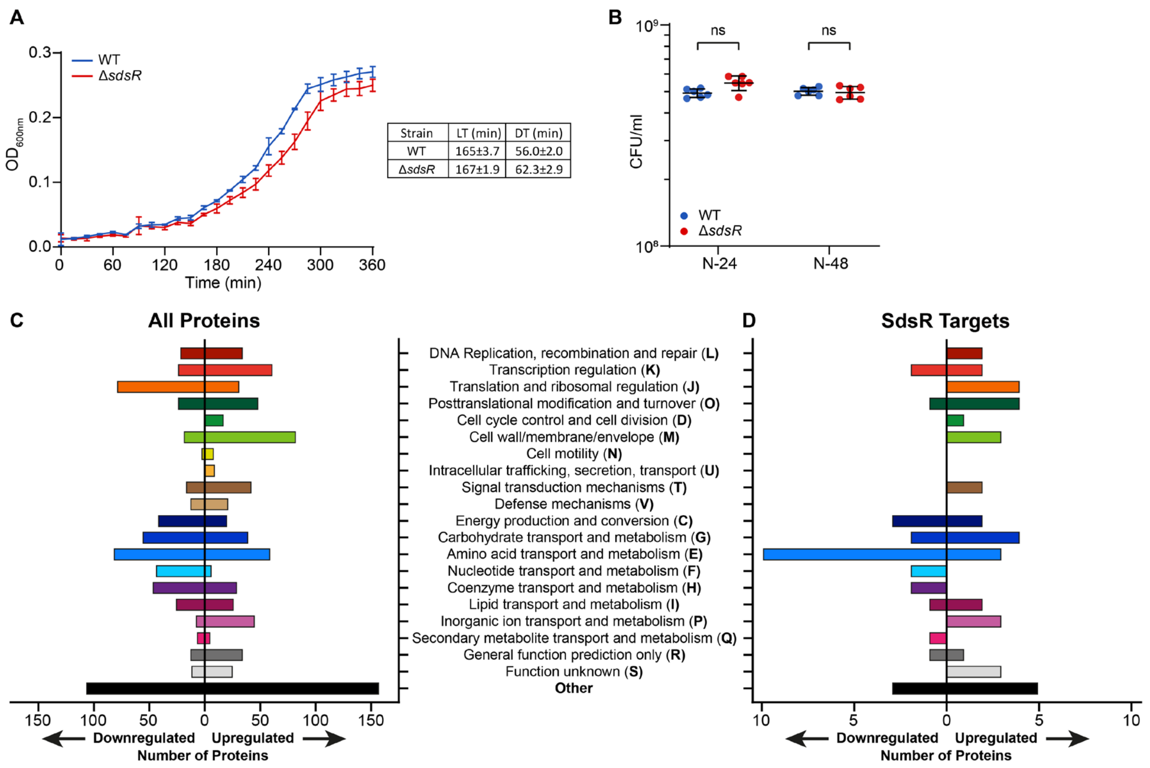


**Figure S5**. **(A)** Growth of WT and Δ*sdsR* bacteria under N limiting conditions. Error bars represent standard deviation (n=3). Tables show lag-time (LT) and doubling-time (DT). **(B)** Viability of WT and Δ*sdsR* bacteria following 24 h (N-24) and 48 h (N-48) of N starvation, measured by CFU counting. Error bars represent standard deviation (n=6). Statistical analysis performed by Welch's T-test. **(C)** Graph categorizing all differentially expressed proteins in Δ*sdsR* bacteria at N-24 by clustering of orthologous groups (COG) annotation**. (D)** As in (C) but only for the targets of SdsR identified by RIL-seq.

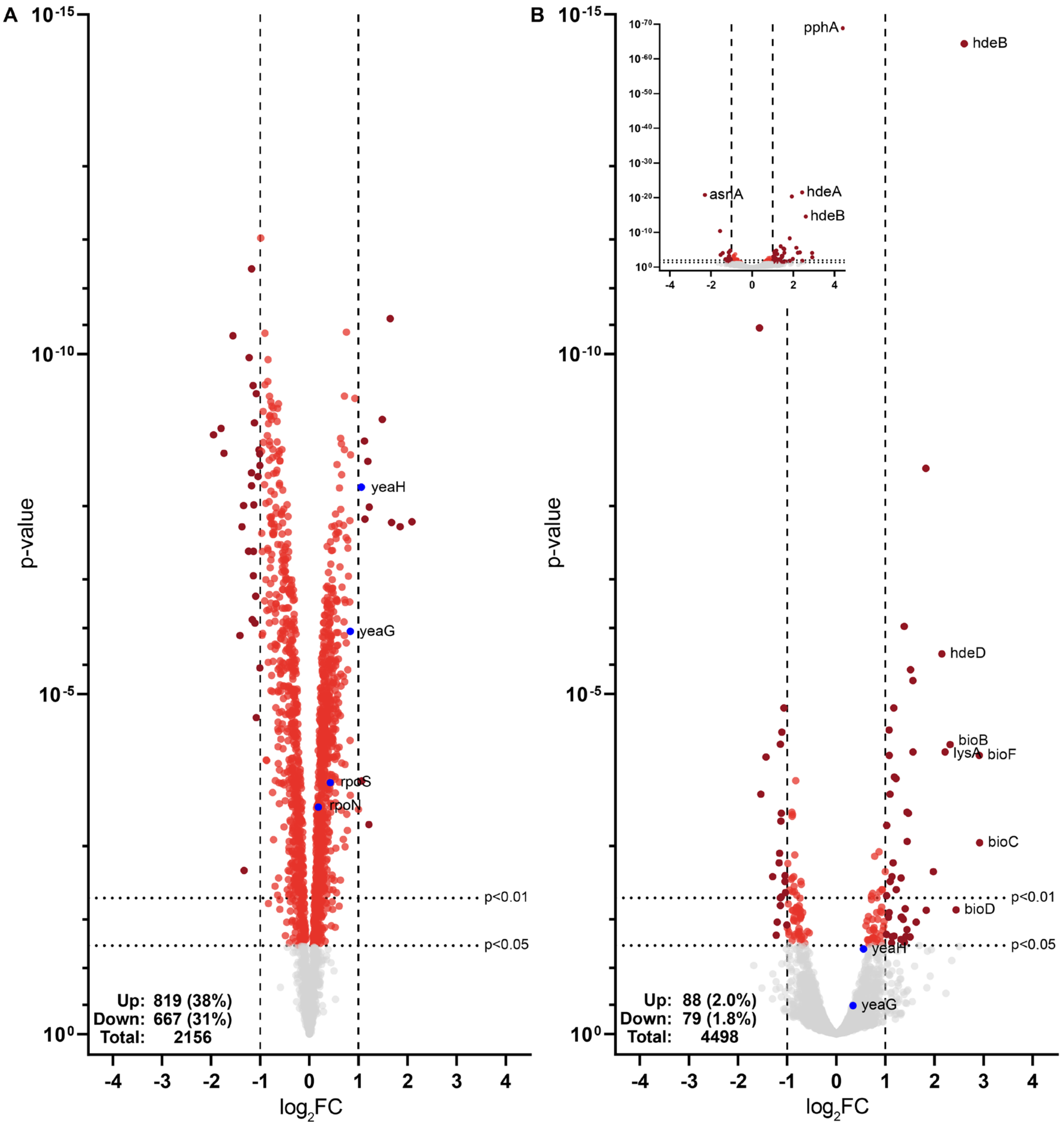


**Figure S6. (A)** Same as Figure 2D. Volcano plot of differential protein levels in N-24 Δ*sdsR* bacteria shown as a $\log_2$ change from wild-type bacteria. Analysis performed by Limma. YeaG, YeaH, RpoS and RpoN are labelled. All proteins differentially expressed with a p-value less than 0.05 are shown in red, with those differentially expressed more than 1 $\log_2$ (i.e., a greater than 2-fold change) are shown in dark red. The number and percentage (of total detected) of differentially expressed proteins are indicated. **(B)** Volcano plot of differential RNA levels in N-24 Δ*sdsR* bacteria shown as a $\log_2$ change from wild-type bacteria. Analysis performed by DESeq2. RNA differentially expressed more than 2 $\log_2$ (i.e., a greater than 4-fold change) and *yeaG* and *yeaH* are labelled. All RNA differentially expressed with a p-value less than 0.05 are shown in red, with those differentially expressed more than 1 $log_2$ (i.e., a greater than 2-fold change) are shown in dark red. Inset was added to allow viewing of genes with very low p-values. The number and percentage (of total detected) of differentially expressed RNA are indicated.

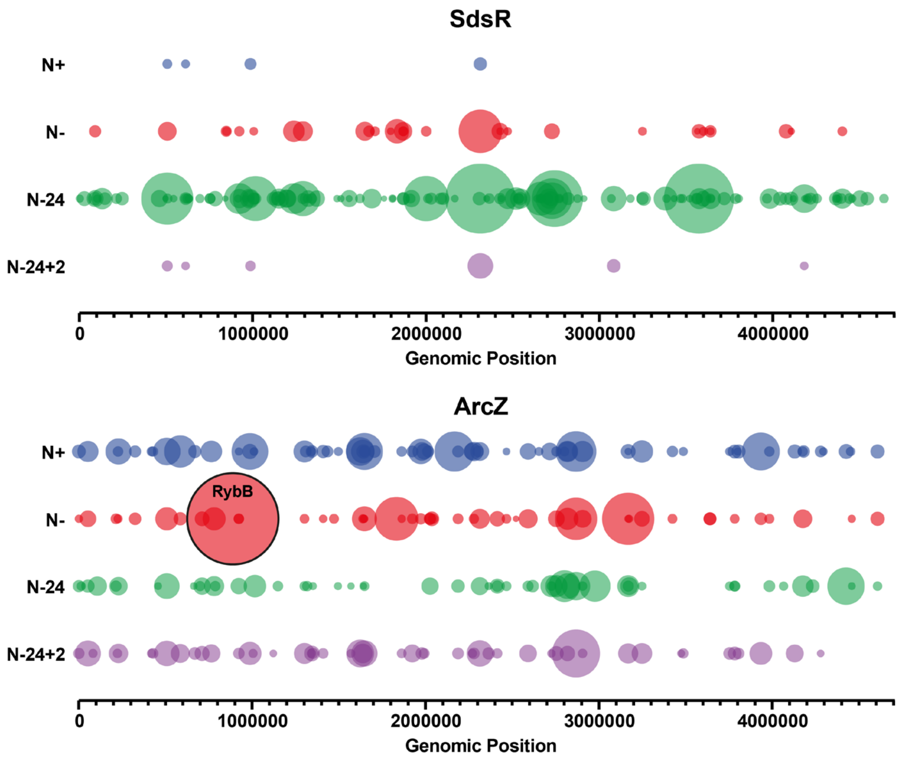


**Figure S7.** Bubble plots of the interaction partners of SdsR and ArcZ at each time point. Each bubble represents an interaction partner of SdsR or ArcZ, at their relative genomic position, scaled such that the area of each bubble is proportional to the number of chimeras that interaction was detected in. RybB is indicated in the ArcZ bubble plot.

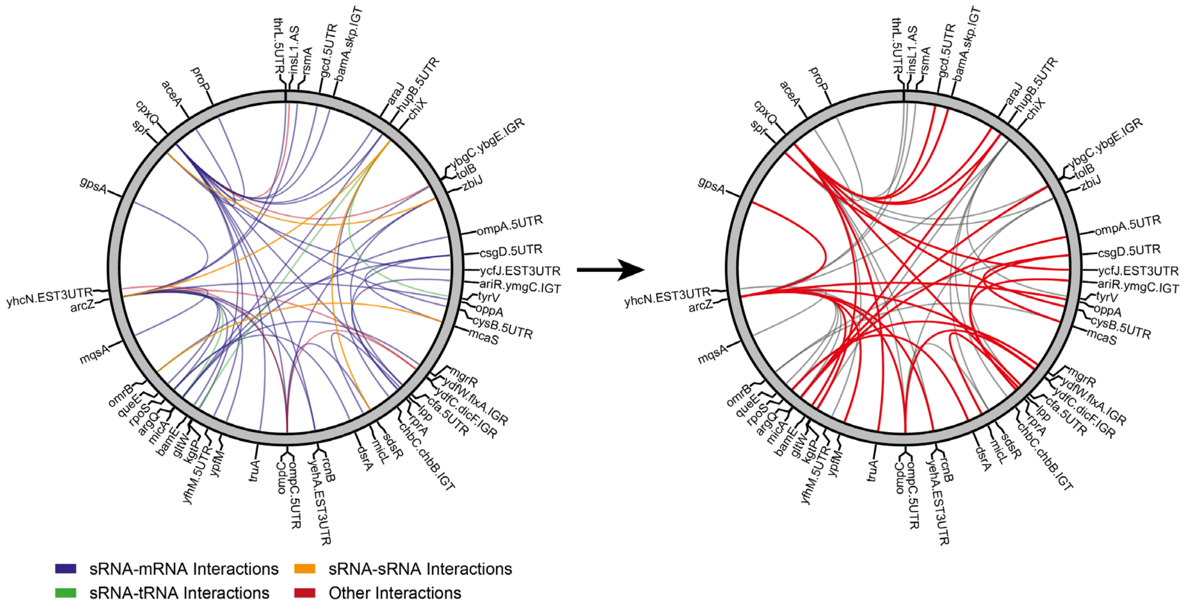


**Figure S8. Left:** Same as Figure 1G. Circos plot of RIL-seq interactions that are detected at in all growth states, represented by at least 30 chimeric fragments in two individual replicates, mapped to the *E. coli* K12 MG1655 genome. sRNA:mRNA, sRNA:sRNA, sRNA:tRNA and other interactions are represented by blue, orange, green and red lines respectively. Thickness of connections are not weighted by the number of chimeras. The RNA involved in each connection are shown. **Right:** Circos plot as in **Left** showing interactions also detected in RIL-seq data from Melamed and colleagues in bacteria experiencing exponential growth and stationary phase in LB, in red (2).

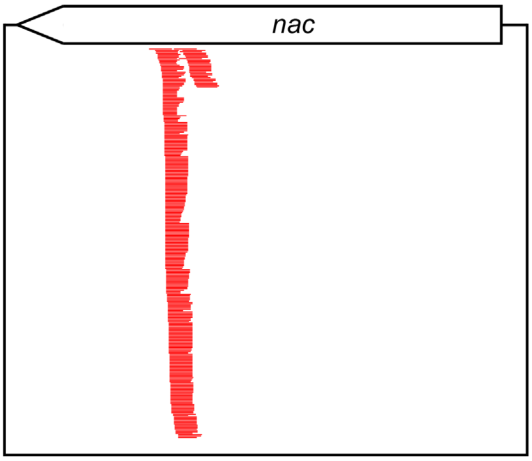


```
nac mRNA   622             632
               |             |
5'-AUG..UUUCC              CACGG..UAA-3'
               CUACGGCGACU
               |:|||||:|||
               GGUGCCGUUGA
3'-UUU..UCAAA              UUUU-5'
               |             |
GlnZ         15              5
```

**Figure S9.** Representative **c**overage plot of chimeras of *glnZ-nac* Chimeric reads are shown in red. Predicted base-pairing interaction site of GlnZ and *nac* mRNA, as determined by IntraRNA (3).

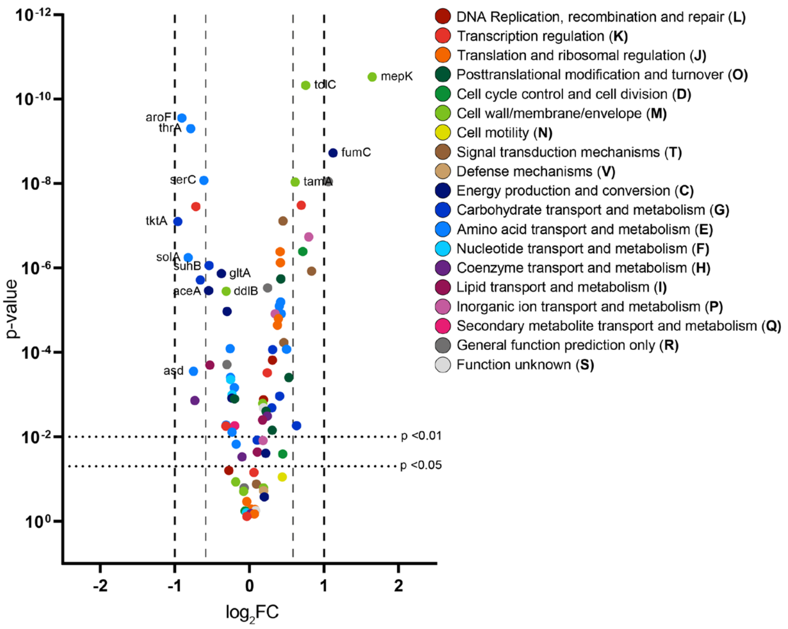


**Figure S10.** Same as in Figure 2D. Volcano plot of differential protein levels in N-24 Δ*sdsR* bacteria shown as a $\log_2$ change from wild-type bacteria, showing only proteins whose mRNA interacted with SdsR as determined through RIL-seq, coloured by clustering of orthologous groups (COG) annotation. Proteins discussed in discussion are labelled.

**Table S1:** Strains and Plasmids used in this study

| **Strains** | | |
|---|---|---|
| Name | Description | Source or Reference |
| Wild-type MG1655 | *E. coli* K-12 *rph*-1 | E. coli Genetic Stock Centre |
| *hfq::*3XFLAG MG1655 | MG1655 *hfq-3XFLAG-kan* | Gift from Prof. Jörg Vogel |
| Δ*sdsR* MG1655 | MG1655 Δ*sdsR::kan* | This Study |
| Wild-type BW25113 | *E. coli* K-12 (*araD-araB*)567 Δ(*rhaD-rhaB*) 568 Δ*lacZ*4787 (*::rrnB-3*) *hsdR*514 *rph*-1 | E. coli Genetic Stock Centre |
| Δ*sdsR* BW25112 | BW25113 Δ*sdsR::kan* | This study |
| Δ*yeaG* BW25112 | BW25113 Δ*yeaG* | (4) |
| Δ*sdsR*Δ*yeaG* BW25112 | BW25113 Δ*sdsR::kan* Δ*yeaG* | This Study(5) |
| **Plasmids** | | |
| Name | Description | Source or Reference |
| pKF68-3 | ColE1 plasmid based on pZE12-luc; expresses *Salmonella* SdsR from $P_{LlacO}$ promoter | (6) |
| pSdsR | pKF68-3 with *Salmonella* SdsR swapped for *E. coli* SdsR | This Study |
| pCONT (pJV300) | Control plasmid, expresses a ~50 nt nonsense transcript derived from *rrnB* terminator | (7) |
| pXG10 | Plasmid backbone to clone translational *gfp* reporter fusions, expresses *gfp* from constitutive $P_{LtetO\text{-}1}$ promoter | (7) |
| p5UTR-*yeaG* | expresses *yeaG*::*gfp* translational fusion (-93 to +63 rel. to AUG) from constitutive $P_{LtetO\text{-}1}$ promoter | This Study |
| p5UTR-*yeaG*$^{MUT}$ | p5UTR-*yeaG* following mutagenesis of residues -28/27 rel. to AUG of *yeaG* from GC to CG | This Study |
| pBR322 | Empty pBR322 | (8) |
| pBR322-*sdsR* | pBR322 expressing *sdsR* from its native promoter, contains 389bp upstream and 248bp downstream of the *sdsR* sequence | This Study |
| pBAD18 | Empty pBAD18 | (9) |
| pBAD18-*yeaG* | pBAD18 expressing 6xHis-*yeaG* under an arabinose-inducible promoter (*araC*) | (4) |
| pBAD18-*yeaG*_K426A | pBAD18-*yeaG* containing a K426A point mutation | (4) |

1. Walling, L.R., Kouse, A.B., Shabalina, S.A., Zhang, H. and Storz, G. (2022) A 3' UTR-derived small RNA connecting nitrogen and carbon metabolism in enteric bacteria. *Nucleic Acids Res*, **50**, 10093-10109.
2. Melamed, S., Peer, A., Faigenbaum-Romm, R., Gatt, Y.E., Reiss, N., Bar, A., Altuvia, Y., Argaman, L. and Margalit, H. (2016) Global Mapping of Small RNA-Target Interactions in Bacteria. *Mol Cell*, **63**, 884-897.
3. Mann, M., Wright, P.R. and Backofen, R. (2017) IntaRNA 2.0: enhanced and customizable prediction of RNA-RNA interactions. *Nucleic Acids Res*, **45**, W435-W439.
4. Figueira, R., Brown, D.R., Ferreira, D., Eldridge, M.J.G., Burchell, L., Pan, Z., Helaine, S. and Wigneshweraraj, S. (2015) Adaptation to sustained nitrogen starvation by Escherichia coli requires the eukaryote-like serine/threonine kinase YeaG. *Scientific reports*, **5**, 17524.
5. McQuail, J., Switzer, A., Burchell, L. and Wigneshweraraj, S. (2020) The RNA-binding protein Hfq assembles into foci-like structures in nitrogen starved Escherichia coli. *The Journal of biological chemistry*, **295**, 12355-12367.
6. Frohlich, K.S., Papenfort, K., Berger, A.A. and Vogel, J. (2012) A conserved RpoS-dependent small RNA controls the synthesis of major porin OmpD. *Nucleic Acids Res*, **40**, 3623-3640.
7. Urban, J.H. and Vogel, J. (2007) Translational control and target recognition by Escherichia coli small RNAs in vivo. *Nucleic Acids Res*, **35**, 1018-1037.
8. Bolivar, F., Rodriguez, R.L., Greene, P.J., Betlach, M.C., Heyneker, H.L., Boyer, H.W., Crosa, J.H. and Falkow, S. (1977) Construction and characterization of new cloning vehicles. II. A multipurpose cloning system. *Gene*, **2**, 95-113.
9. Guzman, L.M., Belin, D., Carson, M.J. and Beckwith, J. (1995) Tight regulation, modulation, and high-level expression by vectors containing the arabinose PBAD promoter. *Journal of bacteriology*, **177**, 4121-4130.
